# Super-omics and PCAS reveal tumor-like molecular landscape of rheumatoid arthritis

**DOI:** 10.1101/2025.01.17.633562

**Authors:** Lanlan Xiao, Nadire Aishan, Siwei Ju, Fengchao Zhao, Xiaoxi Ouyang, Dazhi Chen, Weiqian Chen, Jie Xie, Danhua Zhu, Junchi Zhang, Lingyun Sun, Yimin Li, Dan Zhou, Qingqing Hu, Jinzhi Wu, Qingna Meng, Jichun Zhou, Danyi Xu, Jingyi Yu, Junyu Liang, Yangjun Cai, Minmin Zhang, Yanlin He, Zeyu Sun, Linbo Wang, Jin Lin, Feiyang Ji

**Author notes:** These authors contributed equally to this work (Lanlan Xiao, Nadire Aishan, Siwei Ju, Fengchao Zhao, Xiaoxi Ouyang). **Correspndence:** (F. J.); (J.L.); (L.W.); (Z.S.).

## Abstract

Rheumatoid arthritis (RA) is a debilitating systemic autoimmune disorder that significantly impairs quality of life. To elucidate the molecular alterations in RA progression, we performed comprehensive super-omics analyses on synovial tissues from patients with joint trauma, arthritis, and RA. These analyses included transcriptomics, proteomics, metabolomics, microbiomics, and fourteen PTMs (phosphorylation, acetylation, lactylation, O-GlcNAc glycosylation, arginine monomethylation, lysine monomethylation, lysine dimethylation, lysine trimethylation, succinylation, malonylation, glutarylation, tyrosine nitration, N-glycosylation, and O-glycosylation). Additionally, we developed a protein-centered association study (PCAS) method to integrate these complex datasets. Using this approach, we identified key proteins, such as STK17B, and its interactors, which may be crucial in RA pathogenesis. Tumor-like features, including aberrant angiogenesis and epithelial-mesenchymal transition, were observed in RA, alongside the potential involvement of oncogenes and tumor suppressor genes. Finally, we constructed a molecular interaction blueprint of RA, providing a comprehensive framework for advancing RA pathogenesis, diagnosis, and therapy.

## INTRODUCTION

Rheumatoid arthritis (RA) is one of the most prevalent systemic autoimmune diseases, affecting approximately 0.5% to 1% of the global population^1^. RA is characterized by immune-mediated persistent synovial inflammation, manifesting as symmetric polyarthritis with pain and swelling, typically involving small joints of the hands and feet, and frequently affecting critical organs such as the heart, lungs, and kidneys^2^. The loss of tissue-protective macrophage populations, the emergence of tissue-invasive T effector cells, and the formation of an aggressive fibroblast-like synoviocyte (FLS) phenotype contribute to joint damage^3^. Another hallmark of RA is the production of autoantibodies, particularly anti-citrullinated protein antibodies (ACPA), which are generated through the activation of antigen-presenting cells (APCs) that present citrullinated proteins, further activating specific T and B cells^4^. This leads to the initiation of a series of immune responses that contribute to the autoimmune inflammatory process characteristic of RA. Consequently, RA is widely considered to be an immune and inflammatory disorder. The continuous development and application of biologic and targeted synthetic disease-modifying antirheumatic drugs (b/tsDMARDs) have significantly improved treatment outcomes. However, some patients remain refractory to therapy due to drug resistance, side effects, or other factors, leading to severe impacts on their quality of life. This highlights the need for a more comprehensive understanding of RA.

Advancements in biotechnology, coupled with the development of various omics technologies, have enabled a deeper understanding of the disease. Post-translational modification (PTM) refers to the chemical modification of protein amino acid residues after translation through the addition or removal of specific groups, thereby regulating protein activity, localization, folding, and interactions with other biomolecules. PTMs play a crucial role in various biological processes and disease pathogenesis^5^. Phosphorylation is one of the most common and important PTMs, widely involved in regulating immune cell behavior. Tyrosine nitration is closely associated with cellular oxidative stress^6^, while malonylation is involved in energy metabolism, macrophage inflammatory signaling, and cartilage cell metabolic regulation. Glutarylation has been reported to participate in transcriptional regulation, immune inflammation, and amino acid metabolism ^7^. Lysine and arginine methylation play significant roles in transcriptional regulation and signal transduction^8,9^. Lactylation can affect the function of key immune regulatory molecules, such as NLRP3 and TCR, altering immune cell activity and the intensity of autoimmune responses^10^. Previous studies have shown that succinylation of the mitochondrial transcription factor BRD2 promotes the abnormal differentiation of tissue-resident memory T cells (Trm cells) in RA patients^11^, and acetylation of the T cell microtubule system facilitates T cell migration^12^. In RA patients, the immune response to post-translationally modified proteins is abnormally active^13^, yet proteomic studies on PTMs other than citrullination in RA remain rare.

Existing multi-omics studies have provided valuable insights into the cellular composition and functions of synovial tissues in RA, identifying key factors driving FLS heterogeneity^14,15^, the significance of immune regulation in RA^3,16^, and characteristics of the gut microbiome and serum metabolic profiles in RA patients^17^. However, current multi-omics studies are limited in both the types of omics and the depth of analysis, resulting in incomplete molecular portraits and a lack of specialized data analysis methods for multi-omics data. The analysis of multi-omics data faces significant challenges due to the increasing complexity and diversity of data types. To address these issues, we propose a more advanced multi-omics approach, super-omics, which integrates a comprehensive analysis of key functional molecules in biological activities. Based on this, we have developed a protein-centered association study (PCAS), leveraging known interactions among biomolecules. This new super-omics data analysis pipeline—comprising data preprocessing, differential analysis, PCAS analysis, and network/enrichment analysis—aims to handle the increased omics layers and data diversity, addressing the complexities that arise from traditional analytical methods and identifying key molecular changes.

## RESULTS

### Enhanced molecular insights through super-omics and PCAS

Traditional multi-omics studies^18^ typically combine multiple omics layers, but without strict requirements for the specific levels of omics analysis (such as DNA, DNA modifications, RNA, RNA modifications, proteins, protein modifications, metabolites, etc.). Additionally, these studies often lack sufficient diversity in the area of protein post-translational modifications (PTMs). As a result, both single omics and multi-omics approaches can be limited by the number of omics layers and omics types, leading to outcomes that are often narrowly focused. In these cases, researchers may be so focused on a small number of omics that they can’t see the forest for the trees, failing to provide a comprehensive reflection of the entire biological system and introducing the potential for biased or even inaccurate results. In contrast, super-omics study requires the incorporation of at least ten omics types, spanning at least three omics layers, with a strong emphasis on protein and their PTMs, which are directly involved in the regulation of biological functions. Super-omics offers a significant advancement over traditional multi-omics by broadening both the number of omics types and omics layers, thus offering a more holistic perspective. Furthermore, with advances in mass spectrometry and enrichment method, the detection depth of PTMs has been greatly enhanced, enabling the identification of an expanding array of modifications^19,20^. This progress improves our ability to map the molecular landscape more comprehensively. The Protein-Centered Association Study (PCAS) integrates differential omics data from various layers, based on known molecular interactions and upstream-downstream regulatory pathways, enabling a more nuanced understanding of molecular changes. This approach significantly reduces the risk of error and false discovery rates, ensuring that key alterations are identified (Figure 1).

**Figure 1.**
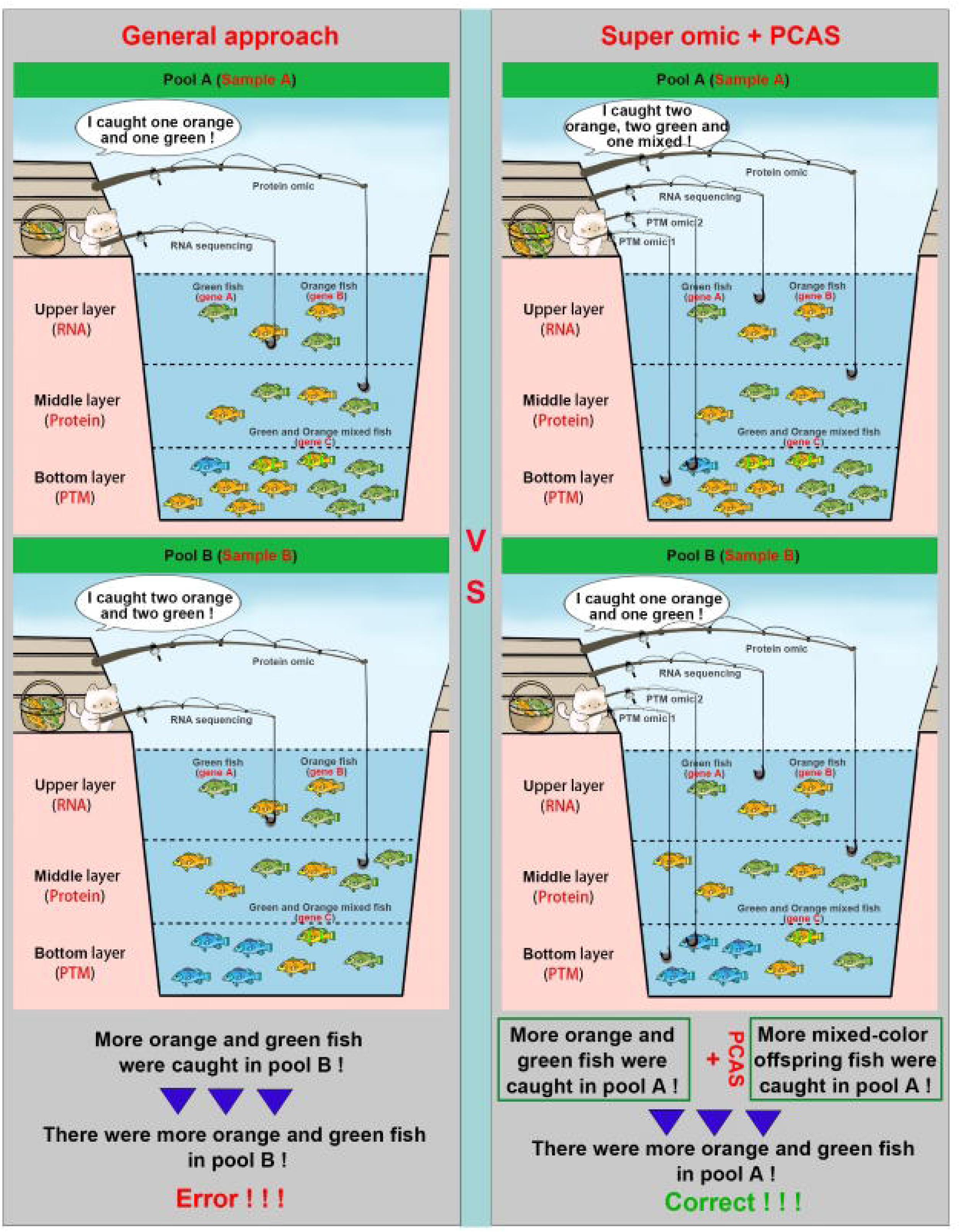
Super-omics and PCAS analytical model. In this analogy, omics can be compared to a fishing rod, and the changes occurring during a biological process are akin to the fish in a pool. The central principle is that changes at different biological levels are represented by fish residing in different layers of the pool. Conventional multi-omics research involves fewer fishing rods and fewer water layers, while super-omics research involves a greater number of rods and more water layers. The PCAS analysis is likened to inferring the number of fish in a pool by examining the parent-child relationships across various fish colors.

### Super-omics characterization of synovial tissue in RA, OA, and NC groups

Between December 2023 and May 2024, we recruited 6 patients with rheumatoid arthritis (RA), 6 with osteoarthritis (OA), and 6 with meniscus injuries (normal controls, NC) (Supplemental Table 1). Synovial tissue samples were collected through Knee Arthroplasty and arthroscopic surgery. We performed a super-omics analysis on these samples, encompassing 18 distinct omics layers: transcriptomics (mRNA), proteomics (WCP), metabolomics (Meta), microbiomics, and 14 types of PTMs, including phosphorylation (P), acetylation (Ac), lactylation (Lac), O-GlcNAc glycosylation (Oglc), arginine monomethylation (Rmme), lysine monomethylation (Pmm), lysine dimethylation (Pmd), lysine trimethylation (Pmt), succinylation (Succ), malonylation (Mal), glutarylation (Glut), tyrosine nitration (Ynit), N-glycosylation (Ng), and O-glycosylation (Og). Our analysis identified 27,459 genes in transcriptomics, 3,424 metabolites in metabolomics, 2,271 amplicon sequence variants (ASV) in microbiomics, and 6,540 proteins in proteomics. PTMs were extensively characterized, with 18,838 phosphorylation sites, 3,420 acetylation sites, 11,684 succinylation sites, 300 lactylation sites, 158 malonylation sites, 74 O-GlcNAc glycosylation sites, 55 lysine monomethylation sites, 69 lysine dimethylation sites, 1,142 lysine trimethylation sites, 70 glutarylation sites, 136 arginine monomethylation sites, 105 tyrosine nitration sites, 566 N-glycosylation sites, and 377 O-glycosylation sites (Figure 2A, Supplementary materials 1). Notably, many proteins displayed multiple PTMs. For example, 440 proteins exhibited both phosphorylation and succinylation, and 144 proteins were modified by acetylation, succinylation, and trimethylation. Among these, alpha-1-antitrypsin (SERPINA1) displayed 10 PTMs, and Mimecan (OGN) displayed 9 PTMs (Figure 2B). The predominant modifications observed were phosphorylation, succinylation, and acetylation (Figure 2C). The correlation analysis of the quantitative results across various omics showed that the within-group correlation in the RA, OA, and NC groups was higher than the between-group correlation, and the within-group correlation among the samples in the RA group was lower than that in the other two groups (Figure S1A). When proteins underwent multiple modifications, the proportion of phosphorylation decreased, while succinylation and acetylation increased, with succinylation emerging as the dominant modification (Figure S1B). The results of the principal component analysis show that in most omics, the RA group is well-distinguished from the OA and NC groups, while the OA and NC groups are closer to each other. At the same time, the samples within the RA group are farther apart, which is consistent with the results of the correlation analysis (Figure S2A). The quantitative results for proteins and PTMs are detailed in Figure S2B (Supplementary materials 2). Among the detected PTMs, O-GlcNAc glycosylation exhibited the highest intensity, followed by phosphorylation, with tyrosine nitration showing the lowest intensity (Figure S2C). N-glycosylation and O-glycosylation were most strongly correlated with protein expression levels (Figure 2D), likely due to their prevalence in secretory proteins^21^. In contrast, arginine monomethylation and lysine monomethylation displayed the weakest correlation with protein expression, potentially because these modifications predominantly occur in nuclear proteins^22^. The relatively low correlation between mRNA and protein levels suggests that protein expression is regulated at post-transcriptional and post-translational levels (Figure 2D).

**Figure 2.**
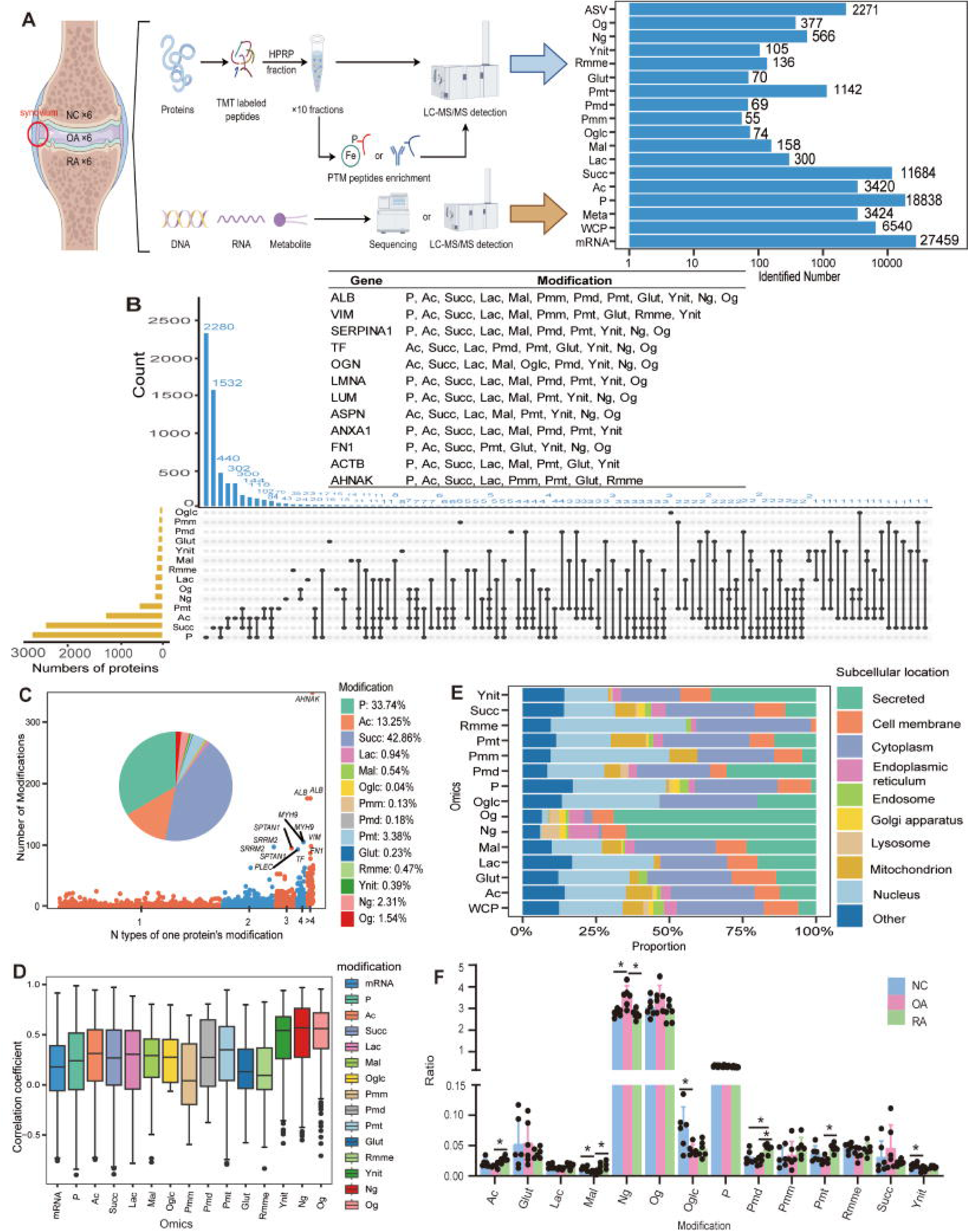
Super-omics analysis of RA synovial tissues. (A) A super-omics analysis on synovial tissue encompassing 18 distinct omics layers and the number of identifications of each omics. RA, rheumatoid arthritis. OA, Osteoarthritis; NC, normal control; TMT, Tandem Mass Tag; LC-MS, Liquid Chromatograph Mass Spectrometer; mRNA, transcriptomics; WCP, proteomics; Meta, metabolomics; ASV, microbiomics; P, phosphorylation, Ac, acetylation; Lac, lactylation; Oglc, O-GlcNAc glycosylation; Rmme, arginine monomethylation; Pmm, lysine monomethylation; Pmd, lysine dimethylation; Pmt, lysine trimethylation; Succ, succinylation; Mal, malonylation; Glut, glutarylation; Ynit, tyrosine nitration; Ng, N-glycosylation; Og, O-glycosylation. (B) UpSet plot shows the distribution of different post-translational modification (PTM) types quantified on the same protein. The table lists the proteins that displayed the most types of PTMs. (C) Scatter plot illustrates the distribution of the number of PTMs per protein, categorized into groups with 1, 2, 3, 4, and more than 4 PTMs. The pie chart represents the proportion of each PTM. (D) Box plot illustrates the correlation between the protein expression level and the transcription level or each PTM level. (E) Stacked bar plot illustrates the subcellular localization profiles of the proteome and each PTM. (F) Scatter-bar plot displays the ratio of the median intensity of each PTM to the median intensity of the corresponding protein across RA, OA and normal NC groups for all detected proteins. * p value < 0.05.

Subcellular localization analysis of omics data revealed that N-glycosylation and O-glycosylation were primarily localized in the endoplasmic reticulum and secretory proteins, indicating their involvement in humoral immunity and inflammatory responses^23^. O-GlcNAc glycosylation was concentrated in the nucleus and cytoplasm, suggesting its role in signal transduction, in line with existing literature^23^. Glutarylation was more prevalent in membrane proteins, implying it may play a role in substance transport, signal transduction, and cell-cell recognition. Acetylation and lysine trimethylation were found more so than others in mitochondria, pointing to their involvement in cellular metabolism^24,25^ (Figure 2E). There are differences in the overall level of certain omics modifications among the three groups. For instance, the levels of acetylation, malonylation, lysine dimethylation, and lysine trimethylation are higher in the RA compared to the OA, while the N-glycosylation level is lower in the RA group compared to the OA group. Additionally, the lysine dimethylation was higher in the RA group compared to the NC group (Figure 2F).

Metabolomics identified 9 major classes of metabolites, with carboxylic acids and derivatives (15.74%) and fatty acyls (13.76%) being the most abundant (Figure S2D). Microbiome analysis revealed that the most abundant bacterial families in the synovial tissue were *Muribaculaceae*, *Prevotellaceae*, and *Lachnospiraceae* (Figure S2E).

### Differential super-omics profiling identifies potential biomarkers and molecular signatures in RA

Comparison of RA with NC revealed significant differences in mRNA, proteins, and PTM sites. Specifically, 1,250 mRNA, 169 proteins, and multiple PTM sites (including 665 phosphorylation sites, 67 acetylation sites, 232 succinylation sites, 12 lactylation sites, 7 malonylation sites, and others) were upregulated in RA, while 1,324 mRNA, 302 proteins, and several PTM sites (including 617 phosphorylation sites and 118 acetylation sites) were downregulated (Figure 3A, Figure S3, Supplementary materials 3). When compared to OA, RA showed upregulation of 381 mRNA, 8 proteins, and several PTM sites, while 1 protein and several PTM sites were downregulated. In entries of the PTMs analyzed in this study, the overall number of upregulated and downregulated PTMs between RA and NC were relatively similar (up: 1133; down: 1204). However, a closer examination of specific PTMs revealed that, compared to NC, the overall levels of acetylation (up: 67; down: 118) and succinylation (up: 232; down: 288) in RA were reduced. When compared to OA, RA exhibited significantly increased levels of phosphorylation and acetylation (Figure S3). Further analysis of the overall differences across various omics showed that the proteomics and phosphorylation differences between the comparison groups were relatively small. Notably, the greatest differences were observed between RA and NC in entries of N-glycosylation, and O-glycosylation, while the overall differences in malonylation were most pronounced between RA and OA, as well as between OA and NC (Figure 3B, Figure S4A). After summarizing the differential entries between all comparison groups across all omics, we found that the intensity variation among the differential entries in the RA group is larger than OA, which may indicate greater synovial heterogeneity in RA patients (Figure 3C). Notably, SYK, and BTK^26^, and protein expression of SUCLG2^12^ were consistent with previous findings in RA research (Figure 3D).

**Figure 3.**
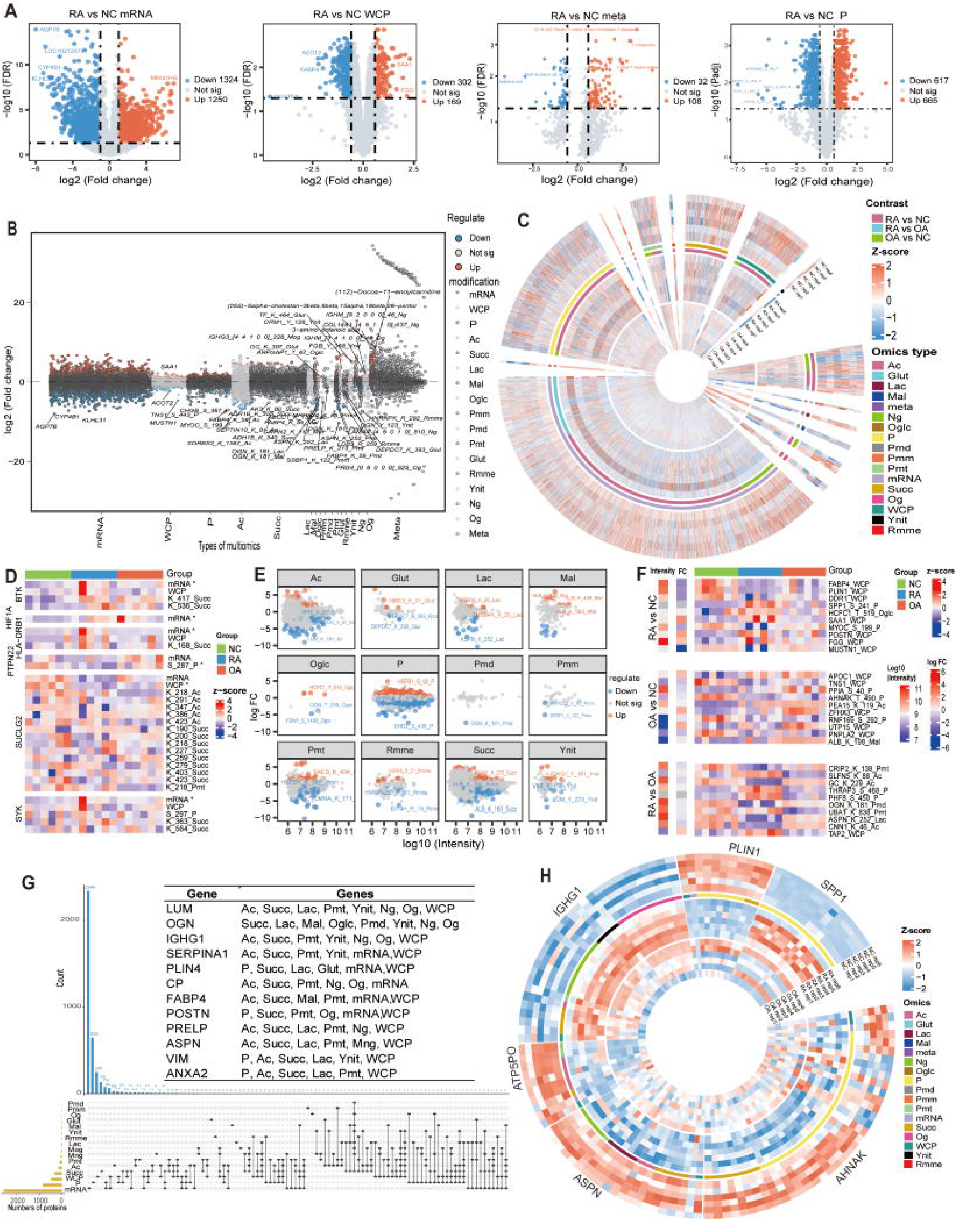
Differential analysis of super-omics data. (A) Volcano plots illustrate differential analysis between RA and NC groups across transcriptomics, proteomics, metabolomics, and phosphoproteomic datasets. The top five most significantly altered features are annotated. (B) The Manhattan plot illustrates the overall differential analysis of transcriptomics, proteomics and 14 types of PTMs between RA and NC groups. Some significantly altered entries are annotated. (C) The circos plot illustrates the intensity of all differential entries across transcriptomics, proteomics and 14 types of PTMs between all comparison groups. (D) The heatmap illustrates the intensity profiles of transcriptomic, proteomic, and statistically significant PTMs associated with known RA-related genes, including HLA-DRB1, PTPN22, ETS1, SUCLG2, HIF1A, SYK, and BTK, across RA, OA and NC groups. * p value < 0.05, which denotes entries exhibiting significant differences between RA and NC groups. (E) The scatter plot illustrates the relationship between the intensity and Fold change (FC) of all detected entries across 12 types of PTMs between RA and NC groups. The top three entries with the highest intensity are annotated. (F) The top five entries with the largest regulation in each omics from the comparison groups were selected. Then the top 10 entries with the highest intensity were chosen. The heatmap illustrates the log (total intensity), log (FC) and z-score intensity for these selected genes. (G) UpSet plot illustrates the distribution of transcriptomics, proteomics, and 14 types of PTMs with differential entries between RA and NC groups on the same gene. The accompanying table illustrates the genes with the highest number of concurrent differential omics. (H) Circos plot illustrates differential entries for genes with the largest number of differential PTMs sites and the largest proportion of differential PTMs sites between RA and NC groups.

In general, greater intensity in PTMs was associated with fewer different between groups, suggesting that housekeeping proteins are more stable and less prone to regulation. However, genes such as ALB, which are typically stable, exhibited significant changes, highlighting their potential role in the pathogenesis of RA. Of course, this could also be due to contamination, as we deliberately retained contaminant proteins when searching the database (Figure 3E, Figure S4B). To screen potential clinical diagnostic markers for RA and OA, we selected the top five entries with the largest regulation in each omics from the comparison groups. After summarizing all omics, the top 10 entries with the highest intensity were chosen. These entries are easier to detect due to their high intensity, and they exhibit better specificity because of the large differences between the comparison groups. For example, serine phosphorylation at position 241 of SPP1 and threonine O-GlcNAc glycosylation at position 519 of HCFC1 can serve as candidate diagnostic markers for differentiating RA and NC (Figure 3F).

In our investigation of differential genes between RA and NC, we identified 20 genes exhibiting concurrent alterations in both phosphorylation and succinylation. Additionally, 13 genes showed changes in succinylation as well as transcriptomic and proteomic alterations, while 4 genes displayed significant differences in succinylation and lysine trimethylation. Mimecan (OGN) was found to be regulated across eight PTMs, including succinylation, lactylation, malonylation, O-GlcNAc glycosylation, lysine dimethylation, tyrosine nitration, N-glycosylation, and O-glycosylation. Lumican showed decreased protein levels, accompanied by downregulation in acetylation, succinylation, lactylation, lysine trimethylation, tyrosine nitration, N-glycosylation, and O-glycosylation, indicating that some of these changes may stem from alterations at the protein expression level (Figure 3G, Figure S4C).

The top 10 genes with the largest number of differential PTMs sites between RA and NC were identified, including AHNAK, IGHG1, ASPN, LAMCA1, LUM, SPTAN1, OGN, ANXA2, TNS1, and IGHM (Figure S4D). Furthermore, genes with the largest proportion of differential PTMs sites were also examined (Figure S4E). Considering both the number and proportion of altered PTMs, we selected genes such as AHNAK, IGHG1, ASPN, ATP5PO, PLIN1, and SPP1 as potential clinical biomarkers for RA (Figure 3H). A joint analysis of transcriptomic and proteomic data revealed 97 genes exhibiting consistent changes in both transcript and protein levels between RA and NC. Among these, SAA1^27^ and GBP5^28^ were notably upregulated at both the transcript and protein levels (Figure S5A), corroborating previous studies that suggest these genes as potential contributors to RA pathogenesis. In contrast, the transcriptional and protein levels of FABP4 and PLIN1 were significantly lower in comparison to NC, with further reductions observed in the phosphorylation and succinylation of PLIN1, and succinylation, acetylation, malonylation and lysine trimethylation of FABP4. Combined analysis of proteomics and PTMs revealed that, compared to the NC group, although no changes were detected in the protein levels of Serotransferrin and SERPINA3 in the RA group, substantial increases were found in the lysine trimethylation and glutarylation of Serotransferrin, as well as acetylation in SERPINA3. OGN exhibited reduced protein expression, with concurrent downregulation in lysine dimethylation and lactylation (Figure S5B).

Metabolic alterations, which are critical for cellular energy homeostasis, were found to be significantly more active in RA compared to both OA and NC, as evidenced by a higher number of upregulated differential metabolites (Figure S5C). Specifically, RA exhibited elevated levels of L-Glutamic acid and D-Lactic acid relative to NC, while sodium lactate was more abundant in RA compared to OA (Figure S5C). Microbiome analysis revealed that the families Muribaculaceae, Rikenellaceae, and Rhodocyclaceae were more prevalent in RA compared to NC, whereas Caulobacteraceae and Beijerinckiaceae were less abundant in RA compared to NC. Additionally, Bifidobacteriaceae and Moraxellaceae were found to be more abundant in RA than in OA (Figure S5D).

### Pathway and kinase analysis reveal immune and inflammatory signatures in RA

To explore the biological functions of RA synovium, we performed Gene Ontology (GO)/Kyoto Encyclopedia of Genes and Genomes (KEGG) enrichment analysis of the significant differential genes from each omics and select the top three entries with the smallest FDR values for each omics. The results revealed enrichment of the “activation of immune response” pathway across eight omics, the “B cell receptor signaling pathway” across nine omics, “lysosomal lumen” across 13 omics, “extracellular matrix structural constituent” across 11 omics, and the “complement and coagulation cascades” across eight omics (Figure 4A, Supplementary materials 4). Besides, differential gene enrichment analysis between RA and OA revealed that the “collagen-containing extracellular matrix” pathway was enriched in seven omics, the “extracellular matrix structural constituent conferring compression resistance” in five omics, and both “vacuolar lumen” and “lysosomal lumen” in four omics (Figure S6A). These findings are in alignment with previous research, highlighting the prominent enrichment of immune and cytoskeletal pathways in RA^29,30^. Further analysis indicated that differential proteins with succinylation were particularly enriched in the tryptophan (Trp) metabolism pathway (Figure 4B). In contrast, the pathways most significantly enriched in the proteomics and phosphorylatomics were thermogenesis and proteoglycans in cancer, respectively (Figure S6B). Recent studies have reported significant changes in Trp metabolites in RA patients, which correlate closely with inflammation, disease activity, and quality of life, and which influence the proliferation and metabolism of fibroblast-like synoviocytes^31^. In our analysis of genes involved in the “activation of immune response” and “rheumatoid arthritis” pathways, we found significant activation of complement and immune pathways in RA, along with high expression of HLA-DRB1 and other human leukocyte antigen (HLA)-specific markers. Elevated mRNA expression of CCL1, CCL2, CCL3, CCL5, CCL3L3, CXCL2, CXCL3, CXCL6, and CXCL12, as well as transcription factors such as IRF1, JUN, MYD88, and TLR1, was observed in RA compared to OA and NC (Figure 4C). Metabolomic analysis highlighted significant enrichment in amino acid metabolism and pathways such as choline metabolism in cancer and central carbon metabolism in cancer (Figure S6C).

**Figure 4.**
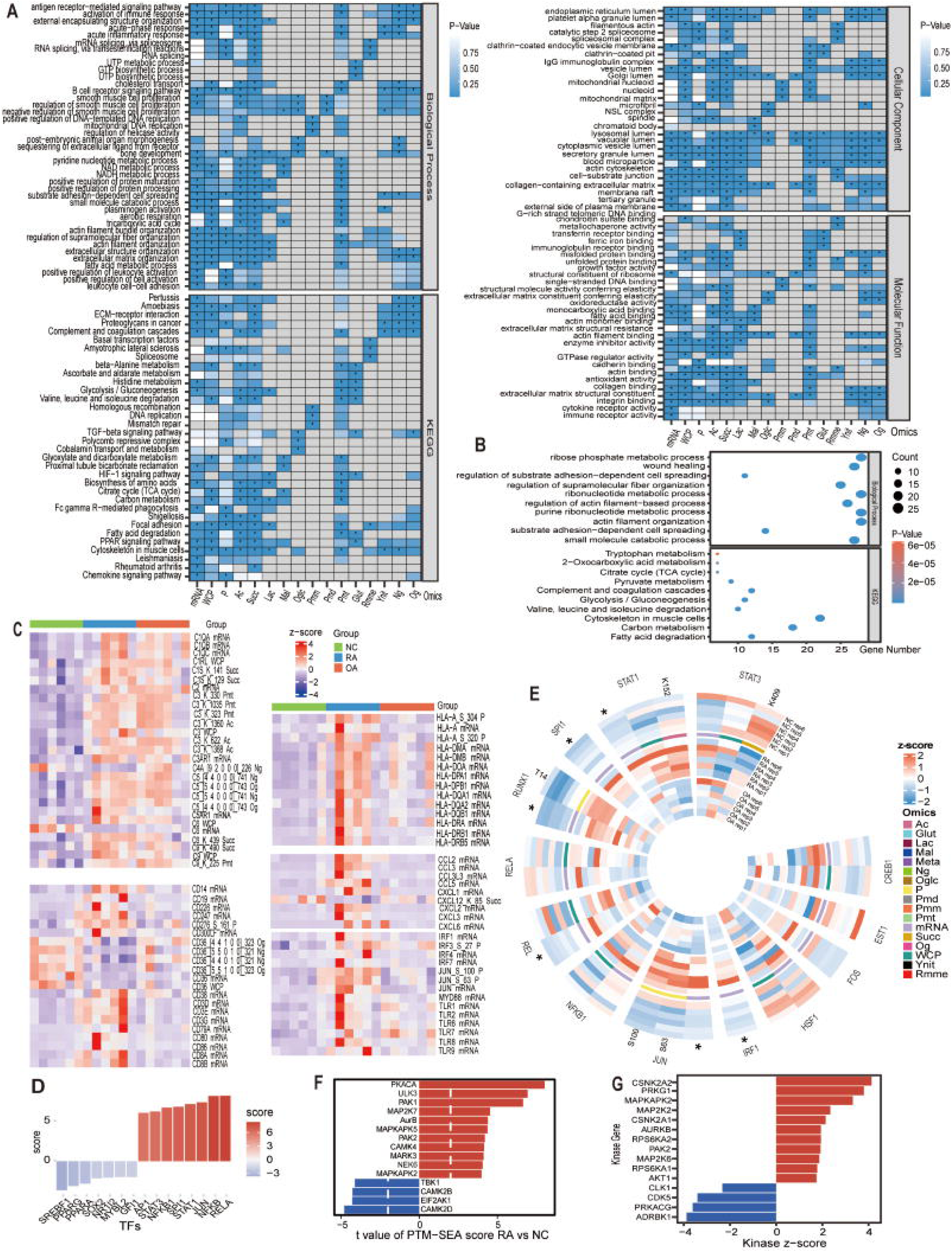
Direct enrichment analysis of differential results from super-omics data. (A) Heatmap illustrate Gene Ontology (GO) /Kyoto Encyclopedia of Genes and Genomes (KEGG) enrichment analysis across transcriptomics, proteomics and 14 types of PTMs between RA and NC groups. *p value < 0.05, which indicates the pathway is significantly enriched. (B) Bubble plot illustrates GO/KEGG enrichment analysis for succinylation between RA and NC groups. (C) Heatmap illustrate the intensity profiles of genes that involved in the “activation of immune response” and “rheumatoid arthritis” pathways across RA, OA and normal NC groups. (D) Bar plot illustrates the predicted inflammation-related transcription factors by comparing the differential gene expression levels between RA and NC groups. (E) Circos plot illustrates the PTM entries of genes from Figure 4D that show differences between RA and NC groups, along with their transcription and protein levels, regardless of whether there is a difference. * p value < 0.05, which indicates genes with differential transcription or protein levels. (F) Bar plot illustrates the top 15 kinases identified by PTM-sea analysis between RA and NC groups. (G) Bar plot illustrates the top 15 kinases identified by KSEA analysis between RA and NC groups.

By comparing the differential gene expression levels between RA and NC, as well as between RA and OA, we predicted that inflammation-related transcription factors such as RELA, NFKB, and JUN are highly active in RA (Figure 4D, Figure S6D). Upon examining the levels of these predicted transcription factors across various omics, we found that, in RA, JUN exhibited higher mRNA and phosphorylation levels compared to both NC and OA. Additionally, STAT1 showed elevated mRNA, protein, and acetylation compared to NC, while STAT3 displayed lower succinylation level in RA relative to NC. RUNX1 also exhibited higher mRNA and phosphorylation levels in RA compared to NC (Figure 4E). PTM-SEA analysis indicated that several inflammation-related kinases, including PKACA, ULK3, and MAP2K7, were activated in RA compared to NC and OA (Figure 4F, Figure S6E, Figure S7A), while KSEA analysis revealed the activation of kinases such as CSNK2A2, PRKG1, CSNK2A1, and MAPKAPK2 in RA (Figure 4G, Figure S7B). These kinases’ phosphorylation profiles showed distinctive patterns, including decreased phosphorylation of RPS6KA1 and PRKG1, and increased phosphorylation of PAK1 in RA compared to NC (Figure S7C). Joint analysis of kinases and their substrates revealed that in the RA and NC comparison groups, the phosphorylation levels of certain substrates were positively correlated with changes in the modification or expression levels of their corresponding kinases. For example, the phosphorylation level of BAD at serine 118 was highly positively correlated with the phosphorylation level of its kinase PAK1 at threonine 103 (R = 0.777, P < 0.05). Similarly, the phosphorylation level of LCP1 at serine 5 was positively correlated with the mRNA level of its kinase RPS6KA1 (R = 0.674, P < 0.05) (Figure S7D). Finally, we plotted the pathways and genes with the most significant changes in RA across all omics into a signaling pathway network diagram, including the activation of immune response, fatty acid metabolic processes, rheumatoid arthritis, extracellular matrix structural constituents, and the PPAR signaling pathway (Figure S7E).

### PCAS analysis identifies key genes and regulatory networks in RA

Super-omics, due to the large number of omics types, leads to a huge number of differentially expressed genes after aggregating all omics data. For example, the comparison between RA and NC groups reveals 3,880 differentially expressed genes (present with differences in at least one omics). Selecting the most critical genes from such a large pool of differentially expressed genes is a significant challenge. The PCAS score integrates various protein-gene, protein-protein and protein-metabolite regulatory relationships to evaluate the significance and reliability of regulated proteins. The score incorporates changes in four dimensions: self, upstream, downstream, and interaction, considering four levels: RNA, WCP, PTM, and metabolic. The PCAS score follows two core principles: (1) Balance between the different omics layers’ ability to reflect protein activity, sequencing depth, and differential analysis algorithm; (2) Enrichment to mitigate errors due to proteins with multiple modification sites, calculating the proportion of differential molecules involved in the same regulatory pathway (Figure 5A). Most importantly, unlike conventional network interaction analysis, PCAS analysis requires the evaluation of molecular relationships in the context of multi-layered omics changes. For instance, the relationship between transcription factors and target genes must satisfy changes in transcription factors at the mRNA, protein, or PTMs levels, as well as changes in target genes at the mRNA or protein level.

**Figure 5.**
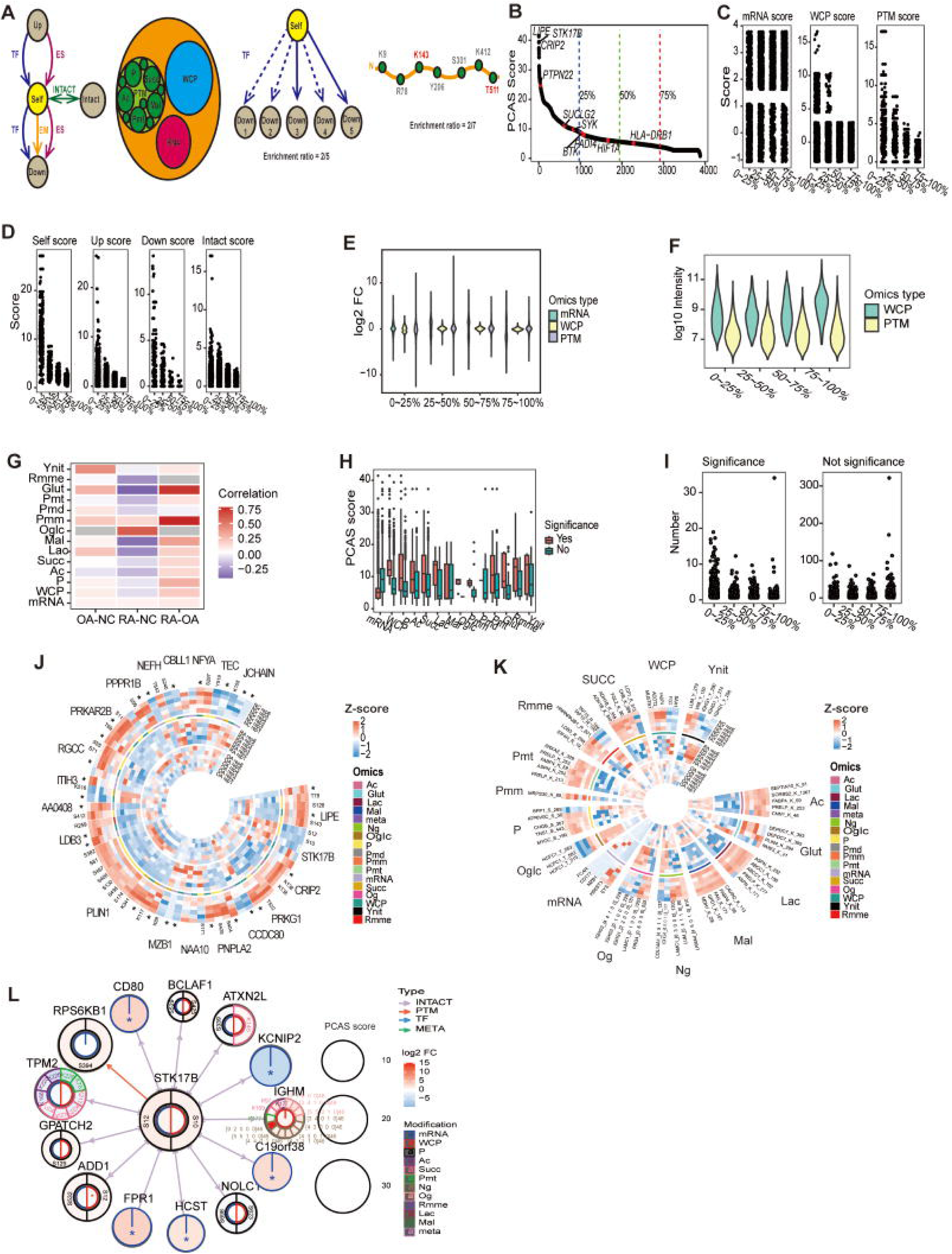
Principles, influencing factors, and analysis results of PCAS. (A) PCAS score incorporates changes in four dimensions: self, upstream, downstream, and interaction; considering four levels: RNA, WCP, PTM, and metabolic; following two core principles: (1) Balance between the different omics layers’ ability to reflect protein activity, sequencing depth, and differential analysis algorithm; (2) Enrichment to mitigate errors due to proteins with multiple modification sites, calculating the proportion of differential molecules involved in the same regulatory pathway. TF, transcription factor-target gene interactions; ES, enzyme-substrate interactions; EM, enzyme-metabolite interactions; INTACT, protein-protein interactions. (B) Scatter plot illustrates the differential genes between RA and NC groups, ranked in descending order by PCAS score. The red dots indicate RA-related genes that have been previously reported. (C) Box plot illustrates RNA, WCP and PTM scores for PCAS score quartile differential genes between RA and NC groups. (D) Box plot illustrates Self, Up, Down and Intact scores for PCAS score quartile differential genes between RA and NC groups. (E) Violin plot illustrates fold change (FC) distribution of PCAS score quartile differential genes across transcriptomics, proteomics and PTM between RA and NC groups. (F) Violin plot illustrates intensity distribution of PCAS score quartile differential genes across proteomics and PTM between RA and NC groups. (G) Heatmap illustrates the correlation between PCAS score and fold change across transcriptomics, proteomics and 12 types of PTM between RA and NC group. (H) Box plot illustrates the PCAS score of genes with significant differences or with non-differential entries across transcriptomics, proteomics and 12 types of PTM between RA and NC group. (I) Box plot illustrates the number of genes with significant differences or with non-differential entries across PCAS score quartile differential genes between RA and NC groups. (J) Circos plot illustrates z-score intensity of top 20 genes based on PCAS score between RA and NC groups across transcriptomics, proteomics and differential PTMs. *p value < 0.05, which indicates differential transcriptomics or proteomics entries. (K) Circos plot illustrates the z-score intensity of 5 genes with the highest absolute values of FC in each omics from top 25% differential genes based on PCAS score between RA and NC groups. (L) The diagram illustrates upstream and downstream relationships of STK17B. It shows the alterations in STK17B phosphorylation and corresponding changes in its upstream and downstream genes.

Upon ranking differential proteins by PCAS score, we found that key RA-related genes, such as PTPN22^26^, SUCLG2^12^, and SYK^32^ were among the top 25% of differential proteins calculated by RA vs NC. Besides, compared with other comparisons, the comparison between RA and NC found more known RA-associated genes (Figure 5B, Figure S8A, Supplementary materials 5). Further analysis of factors influencing the PCAS score revealed that different omics levels and regulatory layers all have an impact on the final PCAS score, indicating that our scoring method indeed integrates multidimensional information (Figure 5C, Figure 5D). The fold change (FC) values in WCP and PTM gradually decreased across the PCAS score quartiles (from 0-25% to 75%-100%), emphasizing the importance of WCP and PTM on PCAS score (Figure 5E). High-intensity proteins were more abundant in the last 25% of PCAS score, indicating greater stability in housekeeping genes (Figure 5F). Notably, the PCAS score of differential proteins from RA and NC showed a weak correlation with FC in tyrosine nitration, but a strong positive correlation with FC in O-GlcNAc glycosylation, and a negative correlation with FC in most omics. In contrast, the PCAS score of differential proteins from RA and OA, as well as OA and NC, exhibited a positive correlation with FC in the majority of omics (Figure 5G). Differential analysis of various omics also revealed that proteins with significant differences in WCP and PTMs, particularly in phosphorylation, acetylation, and succinylation, had higher PCAS scores, whereas transcriptomic data showed the opposite trend (Figure 5H). The top 25% of differential genes by PCAS score between the RA and NC groups exhibited significantly more differential entries compared to differential genes in the other quartiles (Figure 5I). Except for transcriptomics and malonylation, the differential entries in all other omics are primarily distributed among genes with high PCAS scores (Figure S8B). In summary, the PCAS score was derived from an integrative analysis of multiple omics and multiple interaction type.

We further selected the top 20 genes by PCAS score from the differential genes from RA vs NC group, RA vs OA group, and OA vs NC group (Figure 5J, Figure S8C). Taking the RA vs NC group as an example, these genes did not overlap with the top 10 genes identified in single-omics differential analysis between the RA and NC group. Among these genes, we observed that PRKAR2B, PNPLA2, and LIPE exhibited downregulation across transcription, protein, and phosphorylation. PLIN1 showed consistent downregulation in transcription, protein, phosphorylation, and succinylation levels. LDB3 demonstrated downregulation in transcription, protein, and arginine monomethylation, whereas MZB1 and JCHAIN showed upregulation in transcription, protein, and succinylation. CRIP2 exhibited downregulation across protein, lysine trimethylation and acetylation. Remarkably, the directional changes in the transcriptional, protein, and PTMs of these genes were consistent. Additionally, ITIH3 showed downregulation in transcription and upregulation in both protein and succinylation levels, while STK17B displayed no changes in transcription or protein levels, but exhibited increased phosphorylation at S10 and S12 (Figure 5J). All the genes mentioned above and the modifications they undergo may play important roles in the pathogenesis and progression of RA, warranting further investigation to identify new therapeutic targets for RA.

We selected 5 genes with the highest absolute values of FC in each omics from top 25% differential genes based on PCAS score (Figure 5K, Figure S8D). We found that the intensity of several PTMs, such as lactylation, succinylation, malonylation, and lysine trimethylation, in genes obtained from the comparison between RA and NC was generally lower in RA than in NC and OA. In contrast, O-GlcNAc glycosylation exhibited the opposite trend. These differential genes and their corresponding modification changes may serve as potential biomarkers for RA.

The calculation of the PCAS score takes into account changes in four key dimensions: self, upstream, downstream, and interaction. Additionally, it incorporates alterations across four levels of biological data: mRNA, WCP, PTM, and Meta. We present diagrams illustrating the upstream and downstream relationships of top genes based PCAS score from RA vs NC groups, such as STK17B, PLIN1, LDB3, ITIH3, RGCC, and ADH1C (Figure 5L, Figure S9). For instance, changes in the phosphorylation of STK17B lead to downstream alterations in RPS6KB1 phosphorylation, while also interacting with multiple molecules, including TPM2, IGHM, and ATXN2L, all of which undergo various changes of PTMs. Clearly, STK17B undergoes multiple regulatory changes at the molecular, downstream, and interaction levels, which contributes to its high PCAS score. Unlike several other genes, ADH1C is also associated with the differential metabolite Choline (C00114). This analysis highlights the ability of the PCAS analysis method to identify genes with significant regulatory impact.

### RA synovium displays tumor-like molecular landscape

We grouped the top 25% genes based on PCAS score from RA vs NC into several modules according to the interactions among these genes. Modules 1 and 2 were predominantly composed of genes involved in PTM interactions, while Modules 3, 4, 5, 6, and 7 were enriched in genes associated with transcription factor (TF) interactions (Figure 6A, Figure S10A). We then conducted GO/KEGG pathway analyses for the quartile genes based on PCAS score from differential comparison (Figure 6B, Figure S10B). Compared with NC synovium, RA synovium was enriched in immune-related and adhesion-related pathways, in line with the results of the previous individual enrichment analyses of each omics. Notably, RA also displayed enrichment in pathways such as integrin-mediated signaling, SH2 domain binding, and pyruvate metabolism, which were not typically observed in conventional enrichment analyses (Figure 6B, Figure 4A). Gene Set Enrichment Analysis (GSEA), performed on genes ranked by their PCAS scores in descending order, further confirmed the enrichment of RA relative to NC in previously identified pathways, such as kinase activity, focal adhesion, and fatty acid metabolism^12^. Moreover, tumor-related pathways, including epithelial-mesenchymal transition (EMT), VEGFA-VEGFR2 signaling, RAS signaling, transcriptional regulation by TP53, and MYC targets v1, were found to be enriched in RA, highlighting new areas of relevance for the disease. In addition, compared to OA synovium, RA synovium also demonstrated higher enrichment in tumor-related pathways, such as pathways in cancer, cell cycle, and chromatin remodeling (Figure 6C). Furthermore, RA showed significant enrichment in the “PD-L1 expression and PD-1 checkpoint pathway in cancer” compared to OA (Figure S10B).

**Figure 6.**
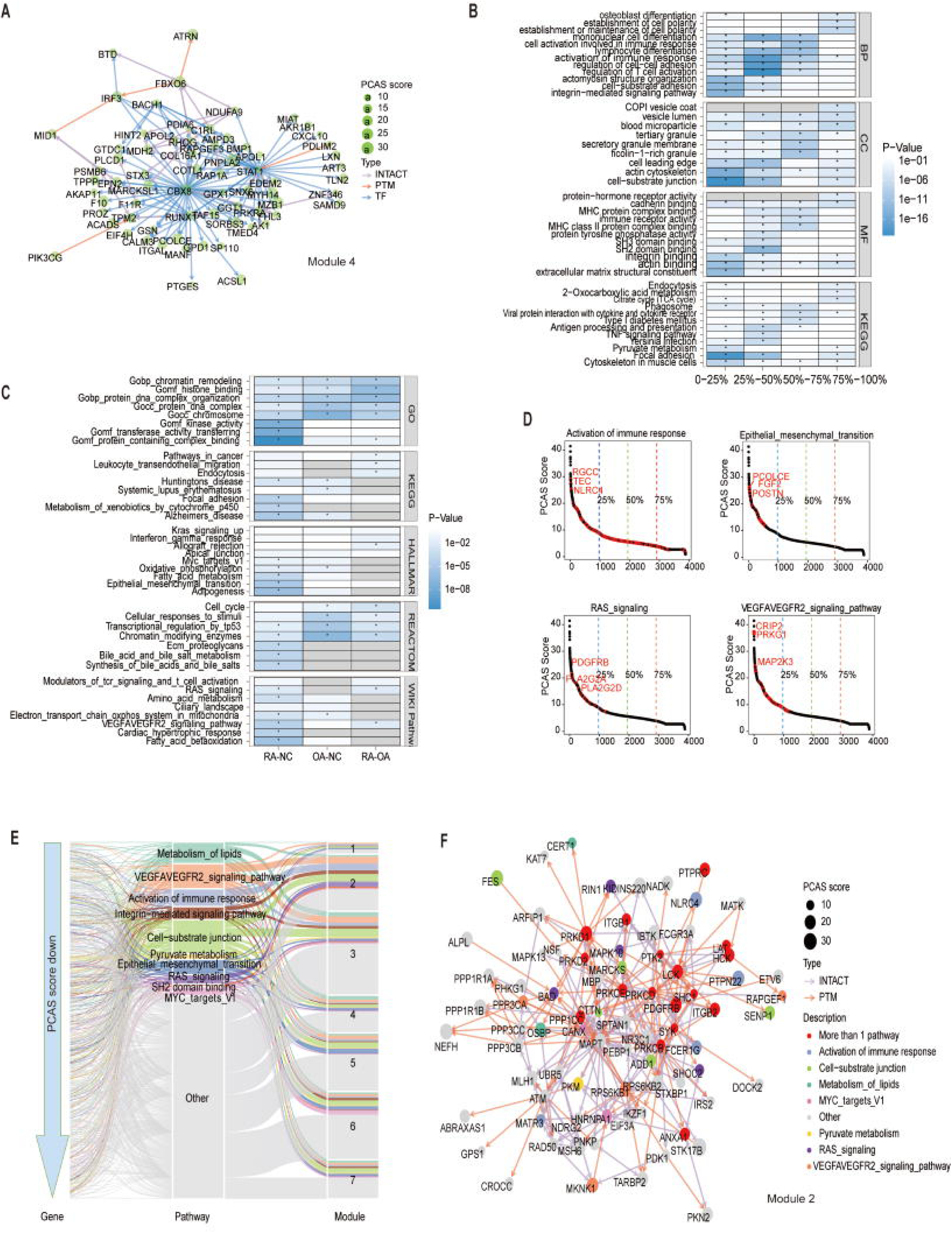
Pathway enrichment analysis and network analysis of PCAS results. (A) Network diagram for Module 4. The Network diagram illustrates three interactions between genes such as INTACT, PTM and TF, as well as the value of the gene’s PCAS score. The top 25% genes based on PCAS score from RA vs NC were grouped into seven modules. (B) Heatmap illustrates the GO/KEGG pathway analysis for quartile genes based on PCAS score between RA and NC groups. (C) Heatmap illustrates the Gene Set Enrichment Analysis (GSEA) analysis performed on genes ranked by their PCAS scores in descending order between RA and NC groups, between RA and OA groups, as well as OA and NC groups. (D) Scatter plots illustrate the genes related with “Activation of Immune Response”, “Epithelial-Mesenchymal Transition”, “RAS Signaling” and “VEGFA-VEGFR2 Signaling” pathways among quartile genes based on PCAS score between RA and NC groups. The red dots indicates the top 3 genes with the highest PCAS scores in the pathway. (E) Sankey diagram integrates the genes from the seven identified modules with the enriched pathways. (F) Network diagram for Module 2 and important pathways. The Network diagram shows pathways for each gene, and genes in Module 2 primarily involve PTMs and protein-protein interactions.

Interestingly, we found that most genes in the epithelial-mesenchymal transition, RAS signaling, VEGFA-VEGFR2 signaling, fatty acid metabolism, MYC targets v1, integrin-mediated signaling, and pyruvate metabolism pathways were enriched among the top 25% of genes based on PCAS score in RA vs NC (Figure 6D, Figure S11A). The integrin signaling pathway, which plays a critical role in signal transduction, angiogenesis, inflammation, and tumorigenesis^33^, and MYC, which can modulate the tumor microenvironment, immune checkpoints, and cytokine secretion to promote tumorigenesis^34^, are both highly relevant to RA. Given that RA is an immune-driven disease with invasive characteristics and that synovial angiogenesis is active, the enrichment of cancer-related pathways in RA synovium suggests that RA shares tumor-like molecular landscape.

Finally, by integrating the genes from the seven identified modules with the enriched pathways, we found that the genes in module 2 had the highest proportion belonging to important pathways such as immune response activation, cell-substrate junctions, and RAS signaling, suggesting that module 2 plays a relatively more important role in RA (Figure 6E). Several top genes based on PCAS score, including STK17B and PTPN22, are also in module 2, and their molecular interactions primarily involve PTMs and protein-protein interactions (Figure 6F). The proportion of important pathway genes in modules 5 and 6 is relatively low, but both modules contain core genes, JUN and FLI1, which interact with almost all other genes (Figure S11B).

### Clinical significance of super-omics and PCAS findings

We conducted an analysis of the top 20 genes based on PCAS score in RA vs NC, and explored their correlations with clinical indicators (Figure 7A). Our findings revealed that the phosphorylation level of STK17B at S12 was positively correlated with erythrocyte sedimentation rate (ESR) and C-reactive protein (CRP) (Figure 7B). Additionally, we performed the same analysis on the most functionally important genes in Figure 6F and found relationships between these genes and clinical indicators, as well. For example, the phosphorylation level of MATR3 at S157 showed a positive correlation with rheumatoid factor and ESR (Figure S11C, Figure 7C). We searched the DGIdb database for FDA-approved drugs and found that the number of FDA-approved drugs targeting the differential genes of RA vs NC decreased progressively across quartiles (from 0-25% to 75%-100%) of the PCAS score (Figure 7D). Besides, we examined the top 20 genes based on PCAS score of RA vs NC, along with genes in Figure 6F, for FDA-approved drugs across the DrugBank, DGIdb, and DSigDB databases (Supplemental Table 2). Notably, several genes within the top 10, including CRIP2, LIPE, PLIN1, PRKG1, and STK17B, had existing targeted drugs.

**Figure 7.**
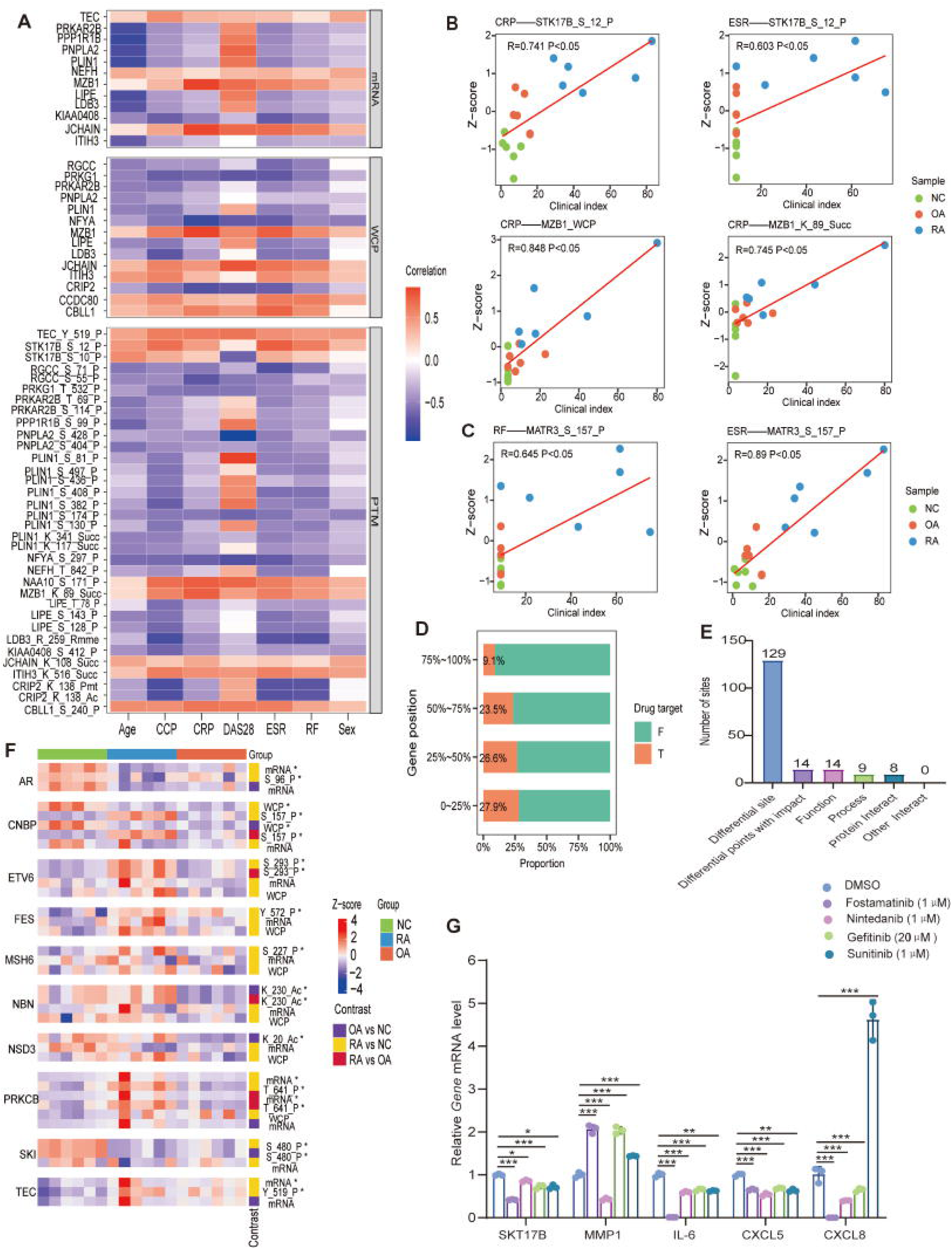
Application research of PCAS results in clinical. (A) Heatmap illustrates the correlation between top 20 genes based on PCAS score between RA and NC groups and clinical indicators. CRP, C-reactive protein; ESR, erythrocyte sedimentation rate; RF, rheumatoid factor; DAS28, Disease Activity Score-28; CCP, Anti-cyclic citrullinated peptide. “SEX” is coded as 1 for male and 0 for female. (B) Scatter plots illustrate the correlation between differential entries of STK17B, MZB1 and clinical indicators (CRP and ESR). (C) Scatter plot illustrates the correlation between differential entries of MART3 and clinical parameters (RF and ESR). (D) The proportion of quartile genes based on PCAS score with FDA-approved drugs targeting the differential genes between RA and NC groups. T, with targeted drug; F, without targeted drug. (E) Bar plot illustrates the number of differential PTM sites, the number of protein associated with known alterations in protein activity and the number of corresponding protein functions. (F) Heatmap illustrates the intensity of the cancer-associated genes within the top 30 genes based on PCAS score of each comparison with differential entries across omics. *p value < 0.05, which indicates entries with significant differences in the corresponding comparison group. (G) The FLSs stimulated with TNFα (10ng/ml), along with Fostamatinib (1μM), Nintedanib (1μM), Gefitinib (20μM), Sunitinib (1μM) for 24h, respectively. *p<0.05, **p<0.01,***p<0.001.

We also assessed the correlation between differential PTMs of these top 20 genes based PCAS score of RA vs NC and their known associations with diseases across OncoKB (Supplemental Table 3). PLIN1 showed high expression in various malignancies. PLIN1 phosphorylation was found to be associated with melanoma and lung adenocarcinoma. Similarly, RGCC phosphorylation was linked to colorectal cancer, while PNPLA2 phosphorylation was associated with neutral lipid storage myopathy. Next, we analyzed the differential PTMs of the top 20 genes based PCAS score of RA vs NC, as well as those in Figure 6F, and examined their effects on protein functional activity (Supplementary Materials 6). A total of 129 differential PTM sites were identified, 14 of which were associated with known alterations in protein activity. These were categorized into 14 functional processes, 9 biological processes, and 8 protein interactions (Figure 7E).

To further explore the connection between RA and cancer, we analyzed the expression of cancer-related genes within the top 30 genes based PCAS score of each comparison. Notably, we found that the transcription and phosphorylation levels of the oncogene AR were lower in RA compared to NC, whereas RA showed higher phosphorylation of the oncogenes CNBP and FES. Additionally, ETV6 and MSH6, both tumor suppressor genes, exhibited increased phosphorylation in RA compared to NC. The acetylation level of NBN was also higher in RA than in OA. On the other hand, the phosphorylation levels of PRKCB and TEC, both cancer-related genes, were higher in RA compared to NC, while SKI exhibited highest phosphorylation levels in NC than in the other two groups (Figure 7F).

To further validate the reliability of the PCAS analysis model, we conducted functional validation of STK17B in FLSs. Following the inhibition of STK17B by siRNA and DRAK2-IN-1 (an STK17B inhibitor), an increase in MMP1 levels was observed, while no significant changes were noted in other inflammatory factors (Figure S11D, Figure S11E). Quercetin, a drug targeting STK17B, effectively inhibits MMP1, IL-6, and CXCL5 (Figure S11E). Previous study^35^ has reported that Quercetin ameliorates RA by reducing joint inflammation, improving joint metabolic homeostasis, and decreasing immune system activation, making it a promising new candidate for RA treatment. Gefitinib, and Sunitinib effectively inhibited STK17B and variably suppressed the levels of IL-6 and CXCL5. However, MMP1 and CXCL8 were not suppressed but rather enhanced under the influence of some of these drugs (Figure 7G). Fostamatinib, Nintedanib, Gefitinib, and Sunitinib are FDA-approved with STK17B as one of their targets. Following intervention with Fostamatinib, Gefitinib, and Sunitinib, MMP1 was upregulated in FLSs, while IL-6 and CXCL5 were downregulated. Nintedanib intervention led to the downregulation of MMP1 and other inflammatory factors (Figure 7G). Fostamatinib^36^, a spleen tyrosine kinase inhibitor, and Nintedanib^37^, a multi-target tyrosine kinase inhibitor, are both used in the treatment of autoimmune diseases. Notably, Nintedanib is specifically used to treat RA-associated interstitial lung disease^38^. These findings suggest their potential therapeutic effects on RA synovium or related pathologies. Gefitinib^39^ and Sunitinib^40^, both anti-cancer drugs, also demonstrated potential therapeutic effects on RA.

In summary, our findings demonstrate that selective inhibition of STK17B leads to upregulation of MMP1 without significant changes in the levels of other inflammatory factors. However, when multi-target drugs, including STK17B as one of their targets, were applied, MMP1 increased while other inflammatory factors decreased. These results suggest that STK17B acts as an anti-inflammatory factor and responds to signaling changes. When other inflammatory pathways are inhibited, STK17B expression is negatively regulated, leading to the upregulation of MMP1.

Through PCAS analysis, we constructed a super-omics landscape, representing the primary genes and crucial pathways involved in the pathogenesis of RA. This landscape not only elucidates the interrelationships between genes, pathways, and their upstream and downstream interactions, but also highlights the potential targeted drugs for each molecular entity, providing a molecular blueprint for future RA research (Figure 8).

**Figure 8.**
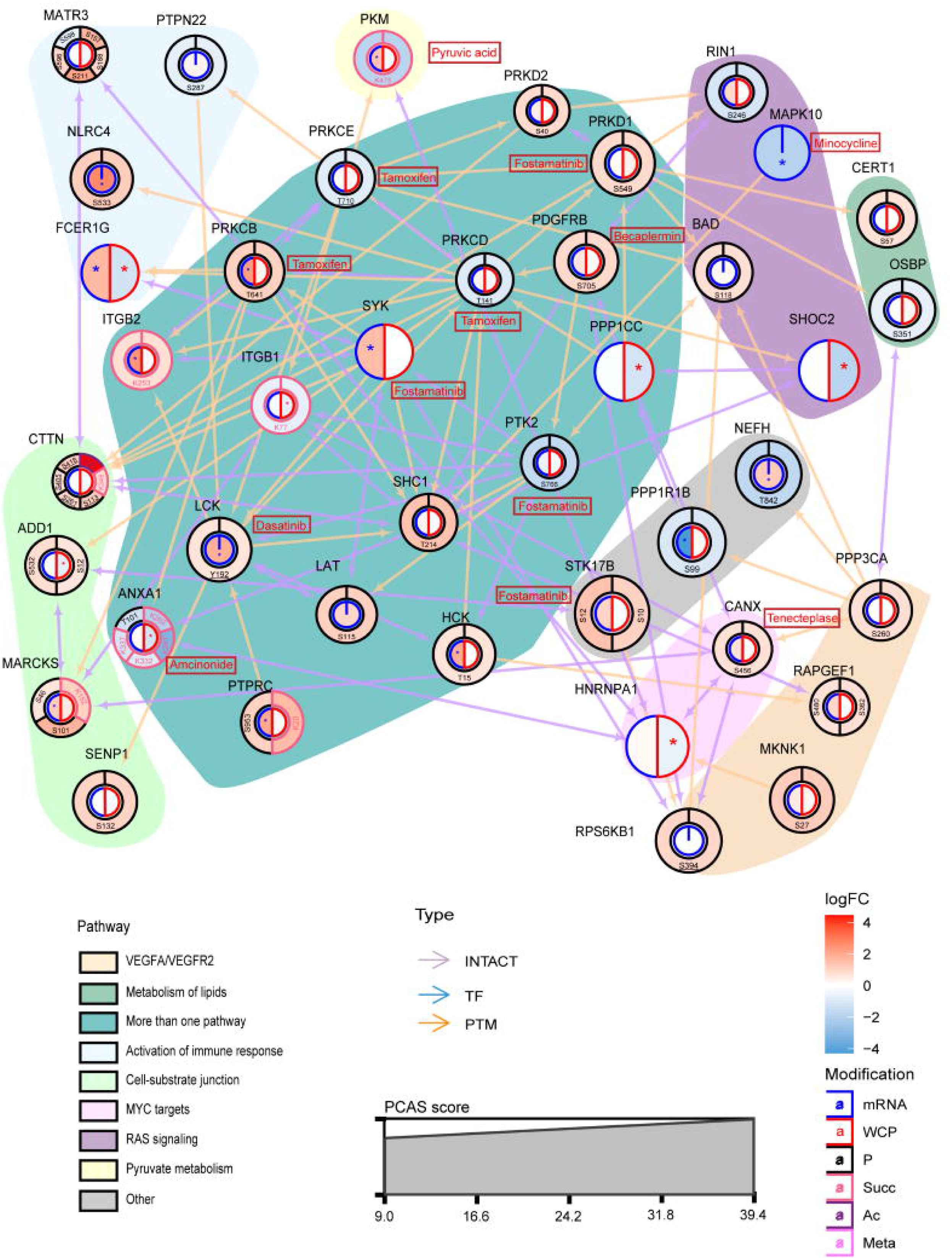
Super-omics landscape of RA. The super-omics landscape elucidates the interrelationships between genes, pathways, and their upstream and downstream interactions, it also illustrates the potential targeted drugs for each molecular entity.

## DISCUSSION

Rheumatoid arthritis (RA) is a systemic autoimmune disorder primarily affecting the synovial joints^41^. The immunopathogenesis of RA spans several decades, beginning with the production of autoantibodies against post-translationally modified proteins. Over the years, this leads to asymptomatic autoimmunity and progressive immune system remodeling, eventually resulting in the transformation of fibroblast-like synoviocytes (FLSs) from benign mesenchymal cells to aggressive, tumor-like cells capable of tissue destruction and invasion^3^. Despite extensive research, the mechanisms by which these cells acquire their pathogenic state remain incompletely understood. Current therapeutic targets for RA include biologic and targeted synthetic DMARDs (b/tsDMARDs), which act on signaling molecules and adaptive immune system components (e.g., TNF-α, IL-6 inhibitors)^42^. Although early inflammation control improves patient outcomes, a significant proportion of patients do not achieve adequate therapeutic goals despite initiating csDMARDs or b/tsDMARDs. Furthermore, some patients, although in remission, face difficulties when tapering their medications. As personalized cancer therapie s continue to evolve, RA precision medicine has lagged behind, and no reliable biomarkers have been established for clinical use^42^. Given the limited understanding of RA’s molecular mechanisms and the paucity of effective treatments, more in-depth research into its underlying pathogenesis is crucial.

In recent years, various omics techniques, particularly the emergence of various post-translational modification (PTM) omics, have provided powerful methods for generating comprehensive molecular profiles of cells and tissues. These approaches offer significant potential for investigating disease mechanisms. However, in RA, systematic multi-omics studies remain scarce, with existing studies often lacking diversity, particularly in the range of PTMs explored. To address this gap, we introduced the concept of super-omics and developed the PCAS method for processing super-omics data. Using these approaches, we conducted a thorough analysis of RA synovial tissue and discovered that RA shares characteristics with tumor-like diseases. We have identified certain genes, such as STK17B, that may play a significant role in the occurrence and development of RA.

Multi-omics refers to the integration of two or more omics, such as genomics, transcriptomics, and proteomics, to conduct comprehensive investigations. The primary goal is to identify composite biomarkers by correlating and deeply integrating data across multiple omic layers. Super-omics, however, extends this approach by integrating over ten different omics, with protein serving as the focal point. In contrast to traditional multi-omics studies, super-omics incorporates at least three omic layers, making it a more robust approach for understanding disease biology.

Multi-omics data provide multi-layered information about biological systems, yet effectively integrating these data to uncover biological significance remains a significant challenge. Traditional multi-omics analyses often rely on differential analysis at a single omics level, followed by statistical correlation and pathway analysis. However, these methods typically integrate only 2-3 omics datasets at a time^43^, limiting their ability to capture the full picture of disease mechanisms. A comprehensive understanding of the underlying system requires joint analysis of multiple data modalities. Recently, the Slobodyanyuk^44^ proposed a novel multi-omics data integration analysis method, with a particular emphasis on directional integration and pathway enrichment analysis. However, directional integration approaches may not fully capture the complexity of biological interactions. Regulatory networks within biological systems are often dynamic and nonlinear, and simplistic linear or hierarchical models may fail to accurately represent the true biological mechanisms, thereby compromising the interpretability of the results. The PCAS analysis method developed in this study integrates data from multiple omics layers, with a focus on functional molecules proteins as the core. By incorporating prior knowledge such as protein-protein interactions, enzyme-substrate relationships, and transcription factor-target gene relationships, it constructs a multidimensional data integration analysis framework. Unlike traditional association analyses based on statistical methods, PCAS integrates known biological knowledge to more accurately identify key molecules and regulatory mechanisms, thereby reducing false-positive results. Additionally, the PCAS method introduces a comprehensive scoring system (PCAS score), which ranks differentially expressed molecules based on three core principles: balance, enrichment, and ambiguity. This helps researchers filter out the most biologically significant key molecules from vast datasets, making the analysis results more concise and intuitive.

Our PCAS analysis revealed several novel findings, particularly regarding cancer-associated pathways that had not been previously highlighted in conventional immune-related analyses. These included the integrin-mediated signaling pathway, VEGFA-VEGFR2 signaling, RAS signaling, MYC targets v1, and SH2 domain binding. These pathways were largely enriched in the top quartile of genes based PCAS score, suggesting their critical importance in RA pathogenesis. Moreover, multiple cancer-related genes within the top 30 genes based PCAS score showed significant changes in phosphorylation and acetylation levels in RA, further supporting the hypothesis that RA may resemble a tumor-like disease. Current studies have substantiated this hypothesis. Clinically, the joint inflammation and destruction in RA patients progressively worse, ultimately leading to joint deformity and functional loss, resembling the local invasiveness of tumors. RA patients often exhibit systemic symptoms such as fatigue, fever, and weight loss, akin to the systemic wasting seen in cancer^45^. RA synovial fibroblasts undergo phenotypic transformation in the inflammatory microenvironment, acquiring tumor-like characteristics such as unlimited proliferation, resistance to apoptosis, and invasive capabilities^46^. The formation of new blood vessels in RA synovial tissue mirrors the angiogenic mechanisms observed in tumors, both representing adaptive responses to inflammatory and hypoxic conditions. The RA pannus provides nutrients and oxygen to the synovial tissue, supporting its persistent proliferation and invasion ^47^. From an energy metabolism perspective, immune cells in RA (T cells, B cells, macrophages, and fibroblast-like synoviocytes) exhibit metabolic reprogramming similar to that of tumor cells. The hypoxic and nutrient-deficient environment in RA synovial tissue promotes metabolic adaptation in immune cells. Metabolic pathways (e.g., glycolysis, oxidative phosphorylation, fatty acid oxidation, and amino acid metabolism) not only supply energy but also regulate cell differentiation, activation, and function, playing a critical role in the initiation and perpetuation of RA^48^. RA tissue shares an inflammatory microenvironment with tumors, characterized by chronic inflammation, immune cell infiltration, and cytokine release^4^. Epigenetic regulation in RA synovial fibroblasts (e.g., DNA methylation, histone modifications) resembles epigenetic alterations in tumor cells, both leading to dysregulated gene expression^49^. Methotrexate and cyclophosphamide are commonly used DMARDs for RA, with methotrexate serving as the anchor drug in RA treatment^50,51^. Notably, methotrexate and azathioprine are also widely used as anticancer agents in clinical practice. They inhibit tumor cell proliferation or induce cell death by blocking nucleic acid synthesis and replication, playing a significant role in cancer therapy^52,53^. RA treatment strategies share similarities with tumor-targeted therapies and immune checkpoint inhibitors, both aiming to block disease progression by inhibiting specific molecular pathways or targets, such as JAK inhibitors^54^, anti-TNF-α agents^55^, and anti-IL-6 drugs^56,57^. The therapeutic outcomes in RA are similar to those in cancer, often manifesting as partial remission and relapse. Even after treatment, some patients may experience recurrent disease^58^. A recent large-scale real-world study^59^ confirmed that RA patients have a higher risk of lymphoma and lung cancer, further suggesting the tumor-like nature of RA.

In conclusion, we have developed and introduced the super-omics approach, which provides a comprehensive molecular blueprint of RA pathogenesis, paving the way for identifying the key genes and pathways involved. While the methodology is still in its infancy, it holds great promise as a robust framework for more integrated and impactful studies of life sciences research. As the saying goes, ‘Don’t miss the big picture while chasing the details’—in the intricate web of molecular mechanisms, focusing on key molecules and their interactions rather than haphazardly dissecting individual functions is vital for advancing RA research. The super-omics landscape presented here plants the seeds for future breakthroughs, offering fertile ground for innovative studies and therapeutic interventions targeting RA.

## ACKNOWLEDGMENTS

We are thankful for the State Key Laboratory for Diagnosis and Treatment of Infectious Diseases. We appreciate the assistance provided by the First Affiliated Hospital of Zhejiang University School of Medicine and the Sir Run Run Shaw Hospital to this study. This study has received support from the National Natural Science Foundation of China (grant numbers 82203610), the National Key R&D Program of China (grant numbers 2022YFC3602003), the Natural Science Foundation of Zhejiang Province (grant numbers LR22H160011), the opening foundation of the State Key Laboratory for Diagnosis and Treatment of Infectious Diseases (grant number SKLID2023KF06), and the National Key Research and Development Program (grant number 2022YFA1303801).

## AUTHOR CONTRIBUTIONS

Conceptualization, X.O., L.W., F.J.; methodology, S.J., Q.M., M.Z., Z.S., F.J.; validation, L.X., S.J., Q.H.; formal analysis, N.A., D.C., J.Z., L.S., D.Z., J.Y., F.J.; data curation, L.X., N.A., F.Z., F.J.; investigation, W.C., J.X., D.Z., Y.L., J.W., J.Z., D.X., J.L., Y.C., Y.H.; writing – original draft, L.X., N.A., S.J., X.O.; writing – review & editing, all authors; supervision, Z.S., L.W., J.L., F.J.; project administration, L.X., N.A., X.O., J.L., F.J.; funding acquisition, J.Z., Z.S., J.L., F.J.;

## DECLARATION OF INTERESTS

The authors declare no competing interests.

## KEY RESOURCES TABLE

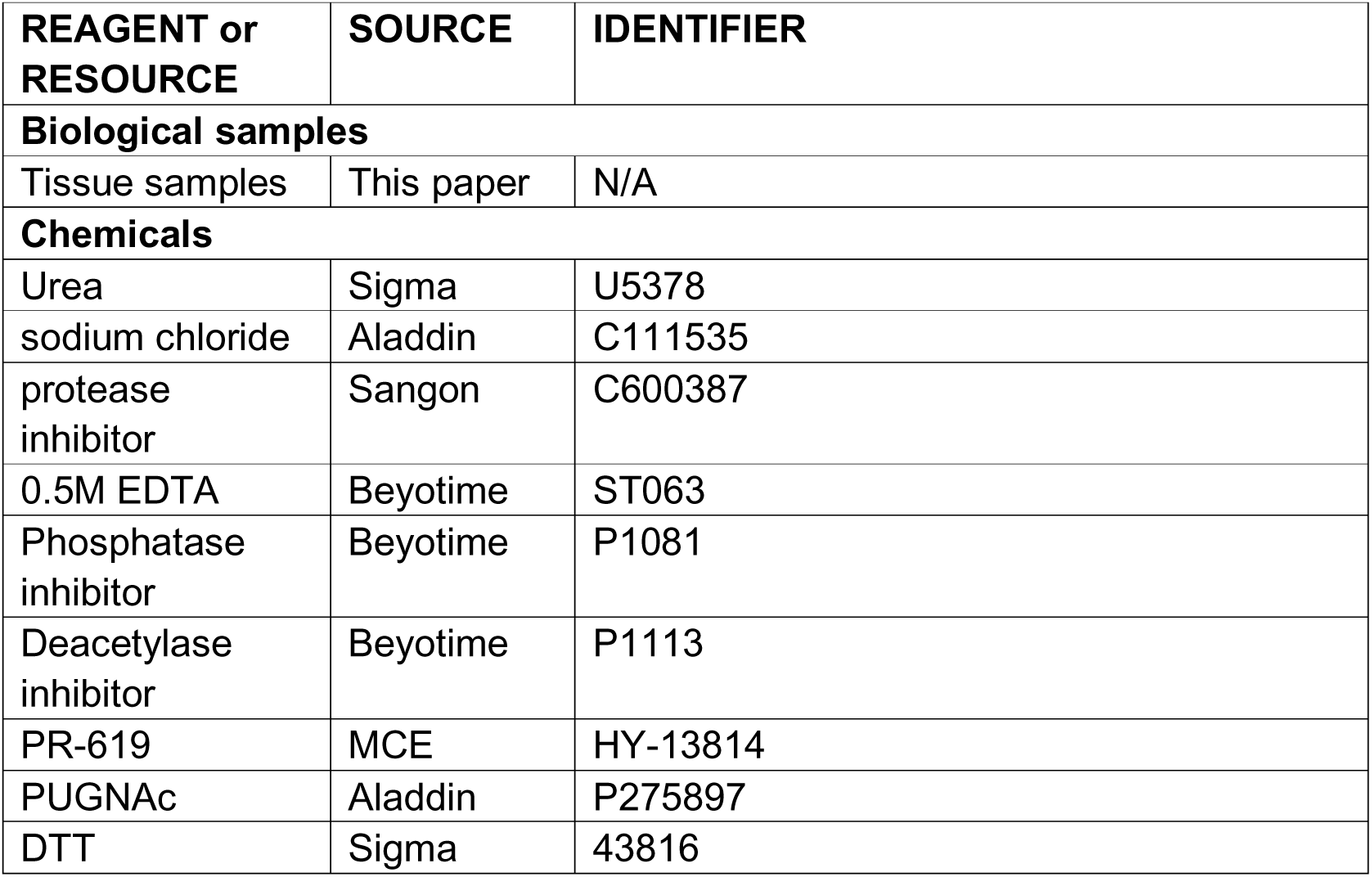
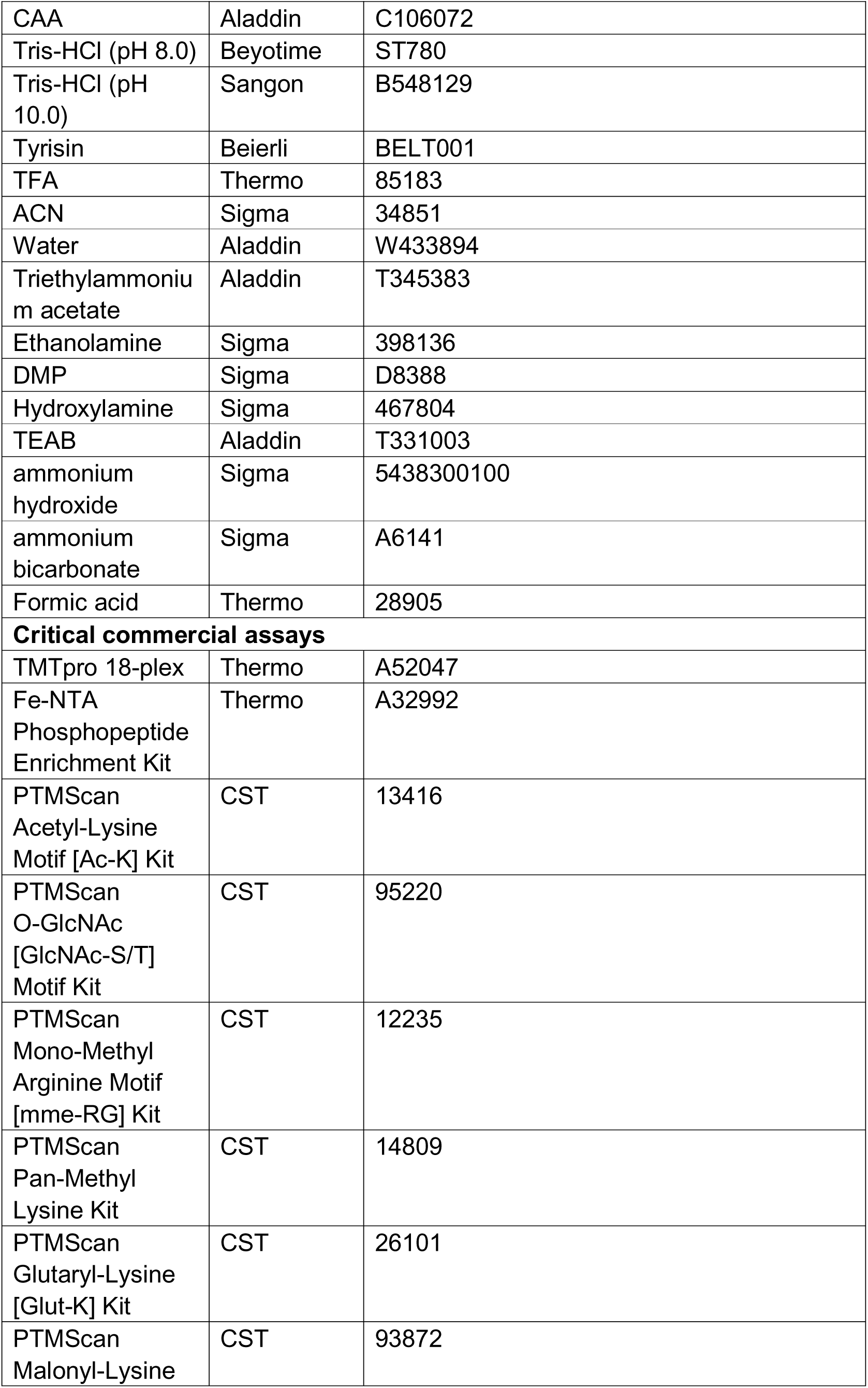
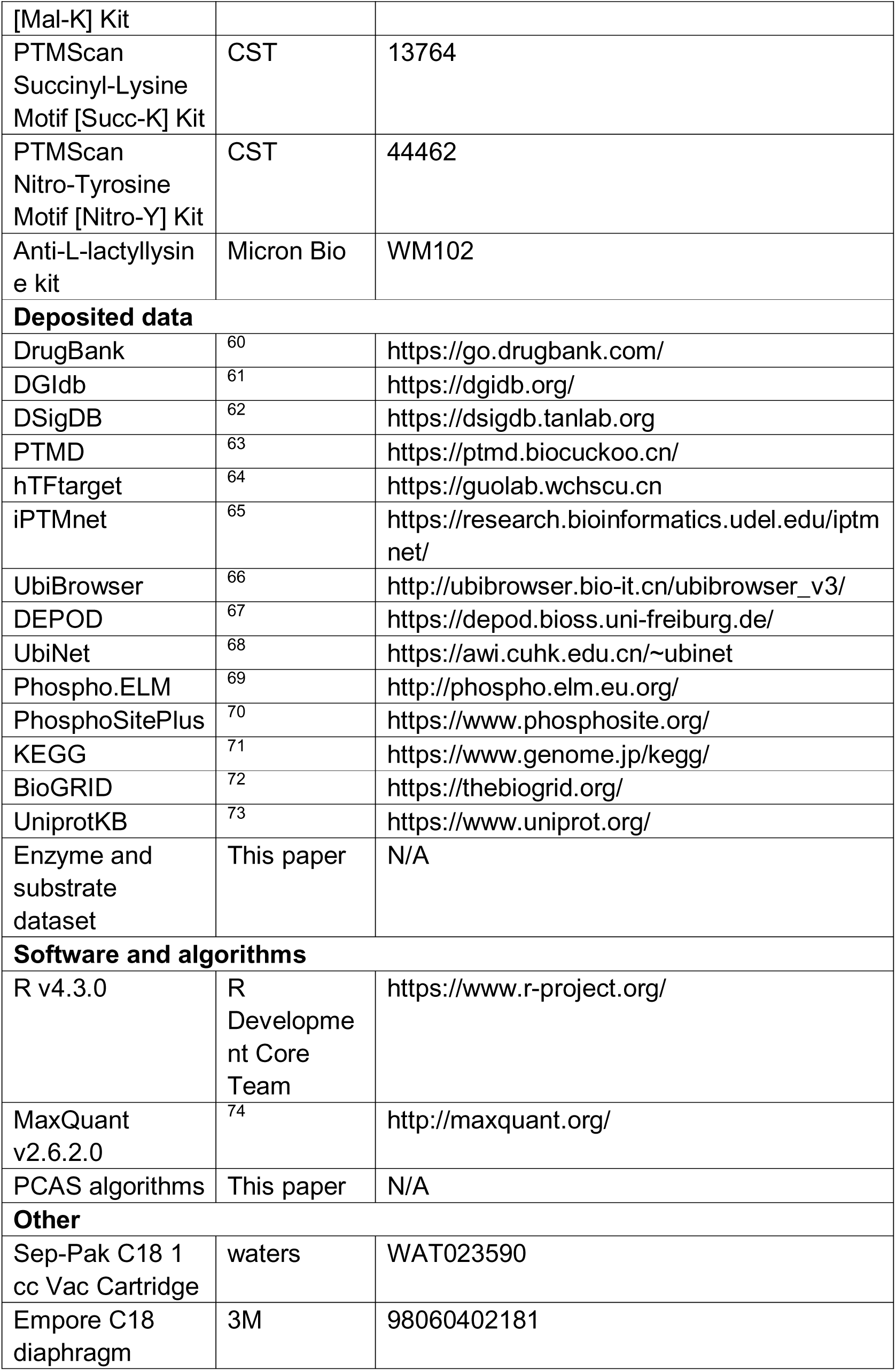

## RESOURCE AVAILABILITY

### Lead contact

Requests for additional information, resources, and reagents should be addressed to the lead contact, Feiyang Ji.

### Materials availability

This study did not generate new unique reagents.

### Data and code availability

Raw proteomic data files and processed proteomic datasets have been deposited in the ProteomeXchange Consortium (https://proteomecentral.proteomexchange.org) via the iProX partner repository (dataset identifier: PXD059446)^75,76^. The raw and processed transcriptomic data have been submitted to the GEO database (dataset identifier: GSE285865)^77^. Raw and processed metabolomic data are available in the MetaboLights database (dataset identifier: MTBLS11642)^78^. All other data can be obtained upon request from the lead contact.

## EXPERIMENTAL MODEL AND SUBJECT DETAILS

### Human Subjects

6 RA patients, 6 OA patients and 6 patients with meniscus injuries (NC patients) were included in this study. RA^79^ and OA^80^ were diagnosed according to the American College of Rheumatology criteria. The detailed clinical information of all Subjects is presented in supplemental Table S1. Ethical approval was obtained from the Ethics Committee of the First Affiliated Hospital, Zhejiang University School of Medicine, and informed consent was obtained from all participants.

## METHOD DETAILS

### Specimen Acquisition

All synovial tissues were obtained from the knee joints of 18 patients who underwent knee replacement surgery and arthroscopic surgery at the First Affiliated Hospital of Zhejiang University School of Medicine. To avoid RNA and protein degradation, all synovial tissues were obtained within 30 minutes after surgery. After washed twice with cold PBS, synovial tissue was immediately freeze in liquid nitrogen after then stored at −80 °C until use.

### Cell purification

RA synovial tissue fragments were incubated with dispase I for 90 min at 37 ℃ and cells were allowed to adhere to tissue culture dishes and passaged every 3-5 days. 3-4 passages yielded a relatively homogeneous population of FLSs.

### Cell Culture

FLSs were cultured in DMEM (plus 10% FBS). The following reagents were used as indicated: TNFα (10ng/ml, PeproTech), DRAK2-IN-1 (1μM, Selleck), Quercetin (80μM, MCE), Fostamatinib (1μM, MCE), Nintedanib (1μM, MCE), Gefitinib (20μM, MCE), Sunitinib (1μM, Selleck). The small interfering RNA (siRNA) targeting STK17B was designed and synthesized by Tsingke Biotechnology Co., Ltd. Lipofectamine RNAiMAX (Thermo Fisher) was used as the transfection reagent, with a final siRNA concentration of 25 nM. Effective inhibition of STK17B was observed in fibroblast-like synoviocytes 24 hours post-transfection.

### Real-time quantitative RT-PCR (qPCR)

RNA was extracted from 0.5×106 FLSs, then reverse transcribed and qPCR was performed.

### 16S Sequencing

#### DNA Extraction and Amplification

Synovial tissue samples were promptly snap-frozen and stored at −80°C following collection. DNA was extracted from the tissues using the MagPure Soil DNA LQ Kit (Qiagen, Guangzhou, China) according to the manufacturer’s instructions. DNA concentration was determined with a NanoDrop 2000 spectrophotometer (Thermo Fisher Scientific, Waltham, MA, USA), and DNA integrity was assessed via agarose gel electrophoresis. The V3-V4 regions of the bacterial 16S rRNA gene were amplified in a 25 µL reaction mixture using universal primers (343F: 5′-TACGGRAGGCAGCAG-3′ and 798R: 5′-AGGGTATCTAATCCT-3′). The reverse primer contained a sample barcode, and both primers were tagged with Illumina sequencing adapters.

#### Library Construction and Sequencing

Amplified products were analyzed by gel electrophoresis to confirm successful amplification. PCR products were then purified using Agencourt AMPure XP beads (Beckman Coulter, USA) and quantified with the Qubit dsDNA Assay Kit (Thermo Fisher Scientific, Waltham, MA, USA). Concentrations were normalized for sequencing. Sequencing was performed on an Illumina NovaSeq 6000 platform, employing paired-end sequencing with 250-base cycles (Illumina Inc., San Diego, CA; OE Biotech Co., Shanghai, China).

#### Bioinformatics Analysis

The raw sequencing data were obtained in FASTQ format. Adapter sequences were trimmed from paired-end reads using Cutadapt software. Following trimming, reads underwent quality filtering, denoising, merging, and chimera removal using DADA2^81^, with default parameters in QIIME2^82^. This process generated representative reads and an Amplicon Sequence Variant (ASV) abundance table. Representative reads for each ASV were selected in QIIME2 and annotated by aligning to the Silva database (Version 138) using the q2-feature-classifier, with default settings. For ITS data, the UNITE database was used for annotation. All 16S rRNA gene amplicon sequencing and subsequent data analysis were carried out by OE Biotech Co., Ltd. (Shanghai, China).

### RNA Sequencing

#### RNA Isolation and Library Preparation

Total RNA was extracted from tissue samples using the RNeasy Lipid Tissue Mini Kit (Qiagen, Hilden, Germany) following the manufacturer’s protocol. RNA purity and concentration were assessed with a NanoDrop 2000 spectrophotometer (Thermo Fisher Scientific, USA), while RNA integrity was evaluated using the Agilent 2100 Bioanalyzer (Agilent Technologies, Santa Clara, CA, USA). RNA-seq library construction was performed using the VAHTS Universal V6 RNA-seq Library Prep Kit as per the manufacturer’s instructions. RNA sequencing and analysis were conducted by OE Biotech Co., Ltd. (Shanghai, China).

#### RNA Sequencing, Quantification, and Normalization

Libraries were sequenced on an Illumina NovaSeq 6000 platform, generating paired-end reads of 150 bp. Approximately 47 million raw reads were generated per sample. Raw reads in FASTQ format were processed using Fastp^83^ to remove low-quality reads, resulting in approximately 46 million clean reads per sample. Clean reads were then aligned to the reference genome using HISAT2^84^. Gene expression was quantified in fragments per kilobase of transcript per million mapped reads (FPKM)^85^, and read counts for each gene were determined using HTSeq-count^86^.

### Non-targeted metabolomics

#### Sample Preparation

Thirty milligrams of the sample were accurately weighed and transferred into a 1.5 mL Eppendorf tube. To this, two small steel beads were added, followed by 400 μL of a methanol-water solution (4:1, v/v) containing a mixed internal standard at a concentration of 4 μg/mL. The mixture was pre-chilled at −40°C for 2 minutes, then homogenized using a bead mill at 60 Hz for 2 minutes. Ultrasonic extraction was performed in an ice-water bath for 10 minutes, after which the sample was incubated at −40°C for 2 hours. The sample was then centrifuged at 13,000 rpm for 20 minutes at 4°C. A 150 μL aliquot of the supernatant was carefully transferred into an LC vial using a syringe and stored at -80°C until analysis by LC-MS. Quality control (QC) samples were prepared by pooling equal volumes of extract from all samples. All reagents were pre-chilled at −20°C before use.

#### LC-MS/MS Analysis

Metabolomic analysis was carried out by Shanghai Luming Biological Technology Co., Ltd. (Shanghai, China) using an ACQUITY UPLC I-Class Plus system (Waters Corporation, Milford, USA) coupled with a Q-Exactive mass spectrometer (Thermo Fisher Scientific, Waltham, MA, USA), equipped with a heated electrospray ionization (ESI) source. Both ESI positive and negative ion modes were employed for metabolic profiling. Chromatographic separation was achieved using an ACQUITY UPLC HSS T3 column (1.8 μm, 2.1 × 100 mm) in both modes. The mobile phase consisted of (A) water with 0.1% formic acid (v/v) and (B) acetonitrile with 0.1% formic acid (v/v). The gradient elution program was as follows: 0 min, 5% B; 2 min, 5% B; 4 min, 30% B; 8 min, 50% B; 10 min, 80% B; 14 min, 100% B; 15 min, 100% B; 15.1 min, 5% B; 16 min, 5% B. The flow rate was 0.35 mL/min, and the column temperature was maintained at 45°C. The injection volume was 3 μL. Mass spectrometric detection was performed over a range of m/z 70 to 1,050 with a resolution of 70,000 for full MS scans and 17,500 for HCD MS/MS scans. Collision energies were set at 10, 20, and 40 eV. The mass spectrometer parameters were as follows: spray voltage, 3800 V (+) and 3000 V (−); sheath gas flow rate, 35 arbitrary units; auxiliary gas flow rate, 8 arbitrary units; capillary temperature, 320°C; aux gas heater temperature, 350°C; S-lens RF level, 50.

#### Data Preprocessing and Statistical Analysis

LC-MS data were processed using Progenesis QI V2.3 software (Nonlinear Dynamics, Newcastle, UK) for baseline correction, peak detection, integration, retention time alignment, and normalization. Parameters included a 5 ppm precursor mass tolerance, 10 ppm product mass tolerance, and a 5% product ion threshold. Compound identification was based on accurate mass-to-charge ratios (m/z), fragmentation patterns, and isotopic distributions, referencing databases such as the Human Metabolome Database (HMDB), Lipidmaps (V2.3), Metlin, and in-house databases. Missing values (>50% of data points missing per group) were imputed by replacing them with half of the minimum detected value. Inaccurate compound identifications (scores <36 out of 60) were excluded. The resulting data matrix was constructed from both positive and negative ion mode data.

Orthogonal Partial Least-Squares Discriminant Analysis (OPLS-DA) was applied to identify metabolites distinguishing the groups. To assess model reliability, 7-fold cross-validation and 200 permutations (Response Permutation Testing, RPT) were performed. Variable Importance in Projection (VIP) scores from the OPLS-DA model were used to prioritize metabolites contributing to group separation. Statistical significance of differences between groups was evaluated using a two-tailed Student’s t-test. Differential metabolites were selected with VIP scores >1.0 and p-values <0.05.

### Liquid chromatography tandem mass spectrometry (LC-MS/MS) proteomics

#### Sample Preparation

Twenty milligrams of joint synovial tissue was homogenized in liquid nitrogen and resuspended in 200 µL of 4°C lysis buffer (8 M urea, 50 mM Tris-HCl pH 8.0, 75 mM NaCl, 2.5 mM DTT, 1 mM EDTA, 1× protease inhibitor cocktail, 1× phosphatase inhibitor cocktail, 1× acetylation inhibitor cocktail, 20 µM PUGNAc). The sample was sonicated in an ice-water bath for 4 min, followed by a 2 min resting period, repeating the cycle three times. The lysate was then centrifuged at 15,000 × g for 20 min at 4°C, and the supernatant was collected.

Protein concentration was determined using the BCA assay. DTT was added to the supernatant to a final concentration of 5 mM and incubated at 30°C for 30 min. After cooling to room temperature, CAA was added to a final concentration of 15 mM, and the sample was incubated at 25°C for 30 min in the dark. Based on the protein concentration, 1 mg of protein from each sample was mixed with 3 volumes of 50 mM Tris-HCl (pH 8.0). Porcine trypsin was added at a 1:50 (w/w) protein-to-enzyme ratio, and the digestion was performed at 25°C for 12 hours with agitation at 200 rpm. TFA was added to the digestion mixture to a final concentration of 0.5%, and the sample was incubated on ice for 10 min. The mixture was then centrifuged at 1500 × g for 10 min at 4°C, and the supernatant was collected. Peptides were desalted using a C18 column, which was washed sequentially with two volumes of acetonitrile (ACN) and three volumes of 0.1% TFA. After acidification, the peptides were passed through the column and wash three times with 0.1% TFA. Finally, peptides were eluted with 50% ACN containing 0.1% TFA and dried under vacuum.

#### Tandem mass tag (TMT) labeling of peptides

The dried peptides were dissolved in 100 µL of 50 mM TEAB (triethylammonium bicarbonate buffer). To prepare the TMT labeling reagent, 40 µL of acetonitrile (ACN) was added to 0.5 mg of the TMT reagent, followed by vortexing and incubation. The dissolved peptides were then transferred to TMT labeling tubes and incubated at room temperature for 1.5 hours. After incubation, 10 µL of 5% hydroxylamine was added to quench the reaction, and the mixture was incubated at room temperature for an additional 15 min. The labeled samples were pooled and dried by vacuum centrifugation. The dried peptides were reconstituted in 2 mL of 0.1% TFA and desalted using a C18 column. After desalting, the peptides were dried again by vacuum centrifugation.

#### Offline bRP fractionation

TMT-labeled peptides were fractionated using an Agilent 1260 chromatography system with solvent A consisting of pure water containing 0.1% triethylamine and solvent B containing acetonitrile with 0.1% triethylamine. The flow rate was set to 0.5 mL/min. Prior to loading the sample, the column was equilibrated for 30 min with the following gradient: 90% solvent B for 10 min, 50% solvent B for 5 min, and 2% solvent B for 15 min. The sample was dissolved in solvent A to a final peptide concentration of 20 mg/mL, followed by centrifugation at 15,000 × g for 5 min at room temperature, and the supernatant was collected for analysis. Fractions were collected every minute, yielding a total of 70 fractions. The liquid chromatography gradient was as follows: 0 min at 2% solvent B, 5 min at 12% solvent B, 54 min at 30% solvent B, 57 min at 36% solvent B, 60 min at 51% solvent B, 65 min at 90% solvent B, and 75 min at 90% solvent B. After collection, the fractions were dried and combined into 10 intervals. One percent of the total peptide was set aside for standard proteomics analysis, while the remaining fractions were used for proteomics analysis focused on post-translational modifications.

#### Phosphopeptide enrichment

The peptides were dissolved in 200 µL of Binding/Wash Buffer (80% acetonitrile with 0.1% TFA) and centrifuged at 15,000 × g for 10 min at room temperature to collect the supernatant. The inner tube of the kit was trimmed at the bottom and placed in a 1.5 mL EP tube, followed by centrifugation at 1000 × g for 30 s at room temperature to discard the filtrate. Next, 300 µL of Binding/Wash Buffer was added to the inner tube, and the centrifugation was repeated at 1000 × g for 30 s to discard the filtrate. This step was repeated once. Afterward, a rubber filter was inserted into the bottom of the inner tube, and the peptide supernatant was added. The top cap was placed on the tube, and the mixture was incubated at room temperature for 30 min, with gentle tapping every 10 min. After removing the rubber filter, the top cap was loosened, and the inner tube was centrifuged at 1000 × g for 30 s at room temperature to collect the filtrate. Binding/Wash Buffer (300 µL) was added to the inner tube, and centrifugation was performed again at 1000 × g for 30 s to discard the filtrate, repeating the step twice. Next, 300 µL of mass spectrometry-grade water was added to the inner tube, followed by centrifugation at 1000 × g for 30 s to discard the filtrate. Finally, the inner tube was transferred to a new 1.5 mL EP tube, and 100 µL of Elution Buffer (5% ammonia solution) was added. The tube was centrifuged at 1000 × g for 30 s to collect the eluate. The elution step was repeated, and both eluates were combined and dried under vacuum.

#### Antibody-based PTM peptide enrichment

To prepare antibody-conjugated beads for the enrichment of post-translationally modified peptides, protein A/G beads were covalently coupled with antibodies. First, the antibody-bound protein A/G beads were centrifuged at 2000 × g for 30 s at 4°C, and the supernatant was discarded. The beads were then washed three times with 1 mL of pre-chilled antibody cross-linking wash buffer (100 mM sodium borate, pH 9.0). Afterward, 1 mL of antibody cross-linking buffer (20 mM DMP in antibody cross-linking wash buffer) was added, and the mixture was incubated on a 3D shaker at room temperature for 30 min. The beads were then centrifuged to remove the supernatant. The beads were blocked by washing twice with 1 mL of pre-chilled antibody blocking buffer (200 mM ethanolamine, pH 8.0), followed by resuspension in 1 mL of antibody blocking buffer and incubation on a 3D shaker at 4°C for 2 hours. The supernatant was discarded after centrifugation. The beads were then washed three times with 1 mL of pre-chilled IAP buffer (50 mM MOPS/NaOH, pH 7.2, 10 mM Na2HPO4, 50 mM NaCl) and stored at 4°C for short-term use. For peptide loading, dried peptides were dissolved in pre-chilled IAP buffer, vortexed, and sonicated. After centrifugation at 15,000 × g for 5 min at 4°C, the supernatant was collected and added to the antibody-conjugated beads. The mixture was incubated on a 3D shaker at 4°C for 2 hours. After centrifugation at 2000 × g for 30 s at 4°C, the supernatant was retained, and the beads were washed five times by adding 0.5 mL of pre-chilled IAP buffer, mixing by inversion, and centrifuging at 2000 × g for 30 s at 4°C. The beads were washed once more with 0.5 mL of pre-chilled mass spectrometry-grade water, following the same centrifugation steps. The beads were then incubated with 50 µL of 0.15% TFA at room temperature on a 3D shaker for 10 min, followed by centrifugation at 2000 × g for 30 s at room temperature to collect the supernatant. This step was repeated, and the supernatants were pooled into a single 1.5 mL EP tube. The collected supernatant was desalted using homemade C18 cartridges and then vacuum-dried.

#### Glycosylated peptide enrichment

The peptides were dissolved in a solution of 95% acetonitrile (ACN) and 1% trifluoroacetic acid (TFA), and centrifuged at 15,000 × g for 5 min at room temperature to collect the supernatant. The MAX column was equilibrated by washing sequentially with 1 mL of ACN (three times), 1 mL of 100 mM triethylammonium acetate buffer (three times), 1 mL of mass spectrometry-grade water (three times), and 1 mL of 95% ACN + 1% TFA solution (three times). The peptide solution was then loaded onto the equilibrated column, and the process was repeated once. After loading the peptides, the column was washed with 0.5 mL of 95% ACN + 1% TFA once, followed by five washes with 1 mL of 95% ACN + 1% TFA. Finally, the glycosylated peptides were eluted with 500 µL of 50% ACN + 0.1% TFA, and the eluate was vacuum-dried.

#### LC-MS/MS analysis

For general proteomics, TMT-labeled peptide samples, prepared after offline bRP fractionation, were desalted, subjected to vacuum evaporation, and then resuspended in 2% ACN and 0.1% formic acid (FA). LC-MS/MS analysis was performed using a Thermo Fisher Scientific Orbitrap Exploris 480 mass spectrometer coupled with an UltiMate NCS-3500RSC system. Gradient elution was carried out at a flow rate of 450 nL/min over 120 minutes. The solvent gradient composition was as follows: 3%-8% solvent B from 0 to 3 minutes, 8%-26% solvent B from 4 to 90 minutes, 26%-38% solvent B from 91 to 110 minutes, 38%-80% solvent B from 111 to 115 minutes, and 80% solvent B from 116 to 120 minutes. MS spectra were acquired at a resolution of 60,000 over a mass range of 400–1200 m/z, with a normalized automatic gain control (AGC) target of 300%. MS2 spectra were collected at a resolution of 15,000, using Turbo TMT and higher-energy collision-induced dissociation (HCD) with a collision energy of 35%. The isolation window was set to 1.6 m/z, and the dynamic exclusion time was 30 seconds.

For phosphoproteomics, the solvent gradient was as follows: 2%-5% solvent B from 0 to 5 minutes, 5%-22% solvent B from 6 to 90 minutes, 22%-35% solvent B from 91 to 110 minutes, 35%-80% solvent B from 111 to 116 minutes, and 80% solvent B from 117 to 120 minutes. The MS spectra were acquired with a resolution of 60,000 over a mass range of 350–1400 m/z, using the same AGC target (300%). MS2 spectra were obtained with a resolution of 15,000, Turbo TMT, and HCD at 35% collision energy, with an isolation window of 1.6 m/z and a dynamic exclusion time of 20 seconds.

For other post-translational modifications (PTMs), the solvent gradient was: 6%-11% solvent B from 0 to 3 minutes, 11%-26% solvent B from 4 to 40 minutes, 26%-40% solvent B from 41 to 52 minutes, 40%-80% solvent B from 53 to 56 minutes, and 80% solvent B from 57 to 60 minutes. MS spectra were recorded at a resolution of 60,000 in the mass range of 350–1400 m/z with a 300% AGC target. MS2 spectra were obtained at a resolution of 15,000, using Turbo TMT and HCD at 35%, with a 1.6 m/z isolation window and a dynamic exclusion time of 20 seconds.

#### Raw data processing

Protein identification and quantification were performed using MaxQuant, except for complex glycosylation analysis. TMT18plex quantification was used for the whole-cell proteome, acetylome, and phosphoproteome. The human UniProtKB database was employed for searching, with automatic reverse database and known contaminants as decoys. Carbamidomethylation of cysteine was set as a fixed modification, while N-terminal acetylation and methionine oxidation were considered variable modifications. For phosphoproteomics, acetylomics, and other post-translational modification analyses, additional variable modifications were included: phosphorylation (79.966331), acetyl-Lysine (42.010565), O-GlcNAc (203.079373), mono-methyl arginine (14.015650), pan-methyl lysine (14.015650; 28.031300; 42.046950), glutaryl-Lysine (114.031694), malonyl-Lysine (86.000394), succinyl-Lysine (100.016044), nitro-Tyrosine (44.985078), and lactyl-Lysine (72.021129). A maximum of five modifications per peptide was allowed, and trypsin was used as the enzyme with up to two missed cleavages. Default MaxQuant settings were used for all other parameters.

For complex glycosylation analysis (including N- and O-glycosylation), raw data were processed using pGlyco and pQuant^87^ for qualitative and quantitative analysis, respectively. pGlyco-N-Human and Multi-Site-O-Glycan were used as glycan databases for N- and O-glycosylation, with TMT18 quantification applied. Max missed cleavages were 2, maximum variable modifications on peptides were 3, and precursor tolerance was 4 ppm. Other settings were kept at the software defaults.

## STATISTICAL ANALYSIS

### Normalization and differential analysis

Transcriptomic data: Sequencing data were quality-controlled, and genes expressed in at least 80% of the samples were retained. Differential expression analysis was performed on the filtered count matrix using the DESeq2 package^88^. Genes with an absolute fold change ≥ 2 and an FDR < 0.05 were considered differentially expressed. p-values were adjusted using the Benjamini-Hochberg method.

Proteomic and PTM data: Raw data obtained from MaxQuant were cleaned by removing reverse hits and data with localization probability < 0.75. Quantitative data were log2-transformed and median-normalized before performing differential analysis using the limma package^89^. A absolute fold change ≥ 1.5 and an FDR < 0.05 were applied to identify significantly differentially expressed proteins.

### Enrichment analysis

Gene Ontology (GO) and Kyoto Encyclopedia of Genes and Genomes (KEGG) Enrichment Analysis: Differentially expressed genes were subjected to GO and KEGG functional enrichment analysis using the clusterProfiler package^90^. This analysis assessed biological processes (BP), molecular functions (MF), cellular components (CC), and signaling pathways related to the genes. A p-value < 0.05 was considered statistically significant.

Gene Set Enrichment Analysis (GSEA): Gene sets from the MSigDB database^91^, including Hallmark, GO, KEGG, Reactome, and WikiPathways, were used for GSEA. ClusterProfiler was utilized to perform GSEA based on gene fold change values or PCAS scores. A p-value < 0.05 was considered statistically significant.

### PTM-SEA enrichment analysis

For phosphorylation modification data, PTM-Signature Enrichment Analysis (PTM-SEA) was performed based on the PTMsigDB v2.0.0 database^92^. Using the ssGSEA 2.0 package^93^, enrichment analysis was conducted to obtain NES scores and p-values from 1000 permutation tests. The analysis identified phosphorylation modification-related features, such as stress disturbances, kinase activities, signaling pathways, and diseases. Differential analysis was used to filter features with a p-value < 0.05 as significantly enriched.

### KSEA kinase activity analysis

Kinase-substrate enrichment analysis (KSEA) was performed on phosphorylation data using the KSEAapp package^94^ to evaluate kinase activities associated with the identified phosphorylation sites. Kinase-substrate relationships were obtained from the PhosphoSitePlus and NetworKIN databases^95^.

### Transcription factor activity analysis

Transcription factor-regulon interactions were retrieved from the CollecTRI database^96^. Transcription factors with at least five target genes were selected. Transcription factor activity was evaluated using a univariate linear model (ULM) based on the fold changes of the target genes.

### Correlation analysis

Pearson correlation analysis was applied to assess the relationships between samples, genes, or genes and clinical indicators. A p-value < 0.05 was considered statistically significant, and the correlation coefficient (r) was calculated to represent the strength of the association.

### PCAS analysis

#### Integration of prior knowledge

Prior knowledge was integrated by compiling relationships from various databases (hTFtarget, iPTMnet, UbiBrowser, DEPOD, UbiNet, Phospho.ELM, PhosphoSitePlus, KEGG, BioGRID, and summarized from the published literature (Supplementary materials 7)), including transcription factor-target gene interactions (TF), enzyme-substrate interactions (ES), enzyme-metabolite interactions (EM), and protein-protein interactions (INTACT), to construct a prior knowledge table. Quantitative and differential gene expression data from transcriptomics, proteomics, PTM, and metabolomic were obtained. The deduplicated quantitative genes from transcriptomics, proteomics, and PTM data were compiled into “GENE”. The deduplicated quantitative genes from transcriptomics and proteomics were compiled into “EXP”, while the deduplicated quantitative genes from proteomics and PTM were compiled into “FUN”. The genes with significant changes were categorized as “GENE_R”, “EXP_R”, and “FUN_R”, respectively, forming the PCAS_ID table.

Using the PCAS_ID table and the integrated prior knowledge table, a PCAS_knowledge table was generated, which was divided into the following sub-tables:

TF Sub-table: Genes in the “GENE” category of PCAS_ID that are transcription factors, and genes in the “EXP” category that are transcription factor target genes.

ES Sub-table: Genes in the “GENE” category of PCAS_ID that are post-translational modification enzymes, and genes in the “FUN” category that are enzyme substrates.

EM Sub-table: Genes in the “GENE” category of PCAS_ID that correspond to enzymes catalyzing metabolites.

INTACT Sub-table: Genes in the “GENE” category of PCAS_ID that are part of protein-protein interactions.

Each sub-table contains a “regulated” column with values of “YES” or “NO”, indicating whether the corresponding gene is regulated (based on “GENE_R”, “EXP_R”, or “FUN_R”). These sub-tables were merged into a comprehensive PCAS_knowledge_use table, which includes columns for the upstream “gene”, downstream “target”, “direction” (values of “YES” or “NO”), “type” (values of TF, ES, EM, or INTACT), and “regulated” status (values of “YES” or “NO”). Duplicate values were removed, and the entries with overlapping relationships between ES and INTACT were excluded.

#### Parameter settings

The following parameters were set based on balance principles: Omics-level Balancing (BL):

BL_RNA_ = 1, BL_WCP_ = 2, BL_PTM_ = 2, BL_META_ = 2

Sequencing Depth (BD):

BD_RNA_ = 1, BD_WCP_ = 1, BD_PTM_ = 1, BD_META_ = 1

Differential Analysis Method (BA):

BA_RNA_ = √(All_RNA_ / Regulated_RNA_), BA_WCP_ = √(All_WCP_ / Regulated_WCP_), BA_PTM_ = √(All_PTM_ / Regulated_PTM_), BA_META_ = √(All_META_ / Regulated_META_)

The following enrichment principles were applied:

Protein Site Enrichment (ES):

ES_RNA_ = 1, ES_WCP_ = 1, ES_META_ = 1, ES_PTM_ = (number of differential sites in the protein / total detected sites in the protein) × (total differential proteins / sum of differential sites in proteins)

Regulatory Mode (ER):

The number of differential molecules for a specific regulation type and direction divided by the total detected molecules of that regulation type and direction.

#### PCAS score calculation

The PCAS score was calculated in three tiers:

1. Omics-Level Score (Scoreomic):
  Score_RNA_ = BL_RNA_ × BD_RNA_ × BA_RNA_ × ES_RNA_
  Score_WCP_ = BL_WCP_ × BD_WCP_ × BA_WCP_ × ES_WCP_
  Score_PTM_ = BL_PTM_ × BD_PTM_ × BA_PTM_ × ES_PTM_
  Score_META_ = BL_META_ × BD_META_ × BA_META_ × ES_META_
2. Regulation type Score (Scoreregulation):
  Score_TF_, Score_ES_, Score_EM_, and Score_INTACT_ were calculated for each regulatory type.
3. Directionality Score (Scoredirection):
  Score_up_, Score_down_, Score_self_, and Score_intact_ were calculated.

The individual scores were computed as follows:
  PACS score = Score_self_ + Score_up_ + Score_down_ + Score_intact_
  Score_self_ = Score_RNA_ + Score_WCP_ + Score_PTM_
  Score_up_ = ER_TF_ × Score_self_ + ER_ES_ × Score_self_
  Score_down_ = ER_TF_ × Score_TF_ + ER_ES_ × Score_ES_ + ER_EM_ × Score_EM_
  Score_intact_ = ER_INTACT_ × Score_INTACT_

For cases where the upstream and downstream genes are the same, an additional coefficient of r = 0.5 was applied. The resulting PCAS scores were sorted in descending order to generate the PCAS score table. The top 25% of proteins based on PCAS score were selected, and clustering was performed using the igraph package. The clustering results were added to the PCAS score table in the “Cluster” column.

## SUPPLEMENTARY FIGURE LEGENDS

**Figure S1.**
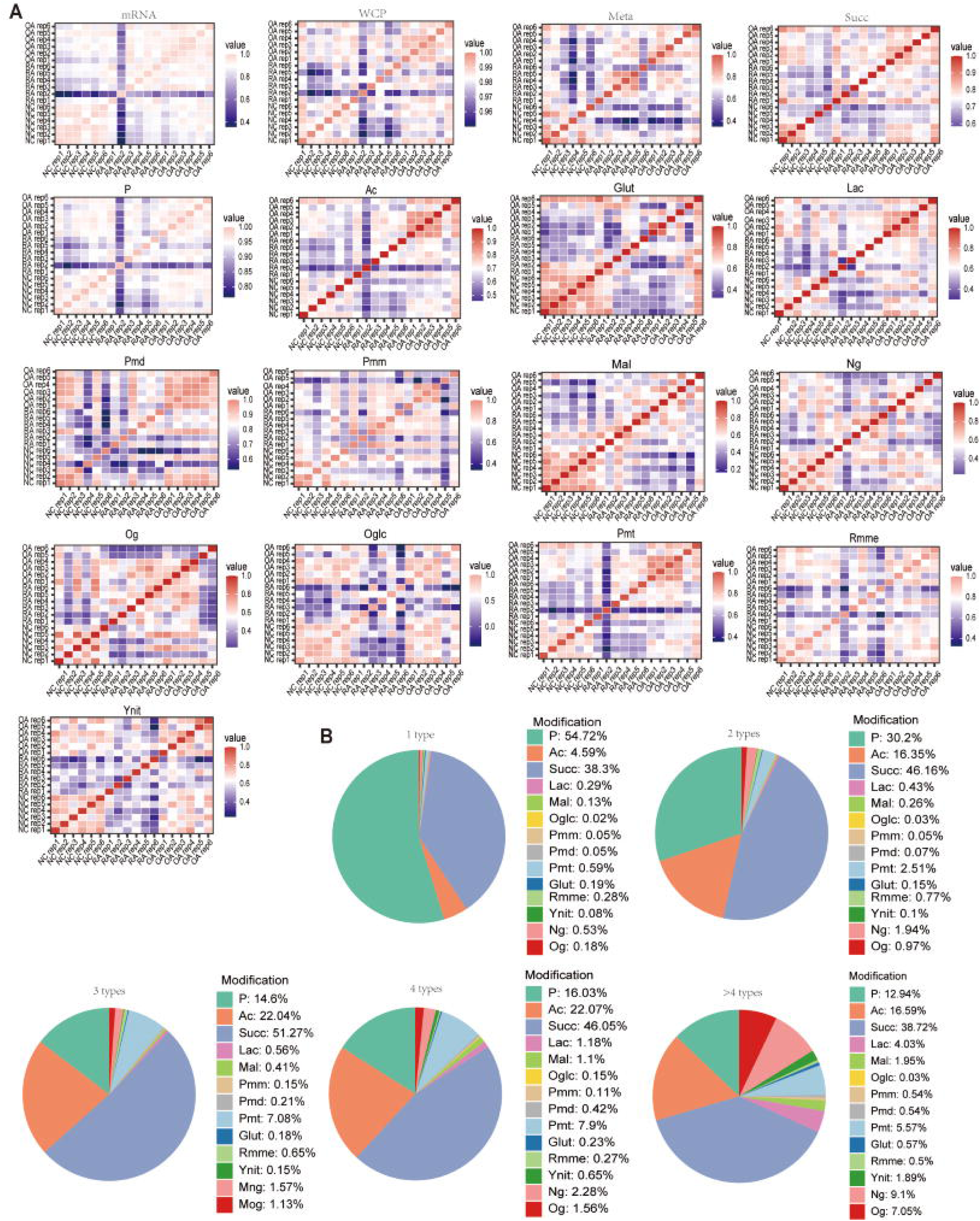
Correlation between different samples in each omics. (A) Heatmap illustrate the correlation analysis of the quantitative results of all samples across various omics. RA, rheumatoid arthritis. OA, Osteoarthritis; NC, normal control; mRNA, transcriptomics; WCP, proteomics; Meta, metabolomics; ASV, microbiomics; P, phosphorylation, Ac, acetylation; Lac,lactylation; Oglc, O-GlcNAc glycosylation; Rmme, arginine monomethylation; Pmm, lysine monomethylation; Pmd, lysine dimethylation; Pmt, lysine trimethylation; Succ, succinylation; Mal, malonylation; Glut, glutarylation; Ynit, tyrosine nitration; Ng, N-glycosylation; Og, O-glycosylation (Og). (B) Pie chart illustrate the proportion of each PTM. 1 type, 2 types, 3 types, 4 types and >4 types indicate the number PTM types of proteins.

**Figure S2.**
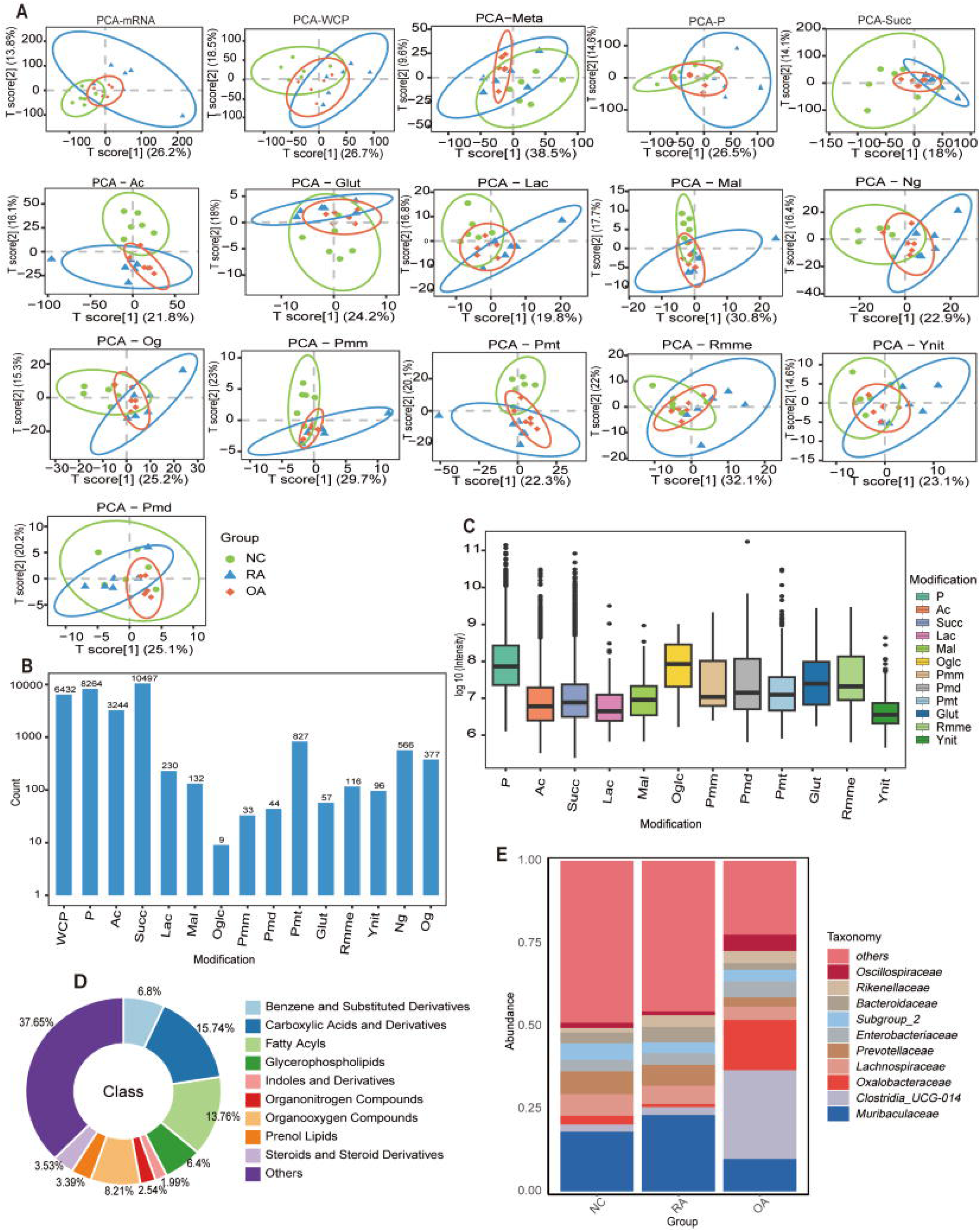
Statistical analysis of qualitative and quantitative identification results of super-omics data. (A) PCA plots of all samples across transcriptomics, proteomics and 14 types of PTMs. (B) Bar plot illustrates the quantitative results across proteomics and 14 types of PTMs. (C) Box plot illustrates the total intensity values of all samples across 14 types of PTMs. (D) Pie chart illustrates the proportion of identified major classes of metabolites in all samples. (E) Stacked bar plot illustrates the relative abundance of bacterial families in RA, OA and NC groups.

**Figure S3.**
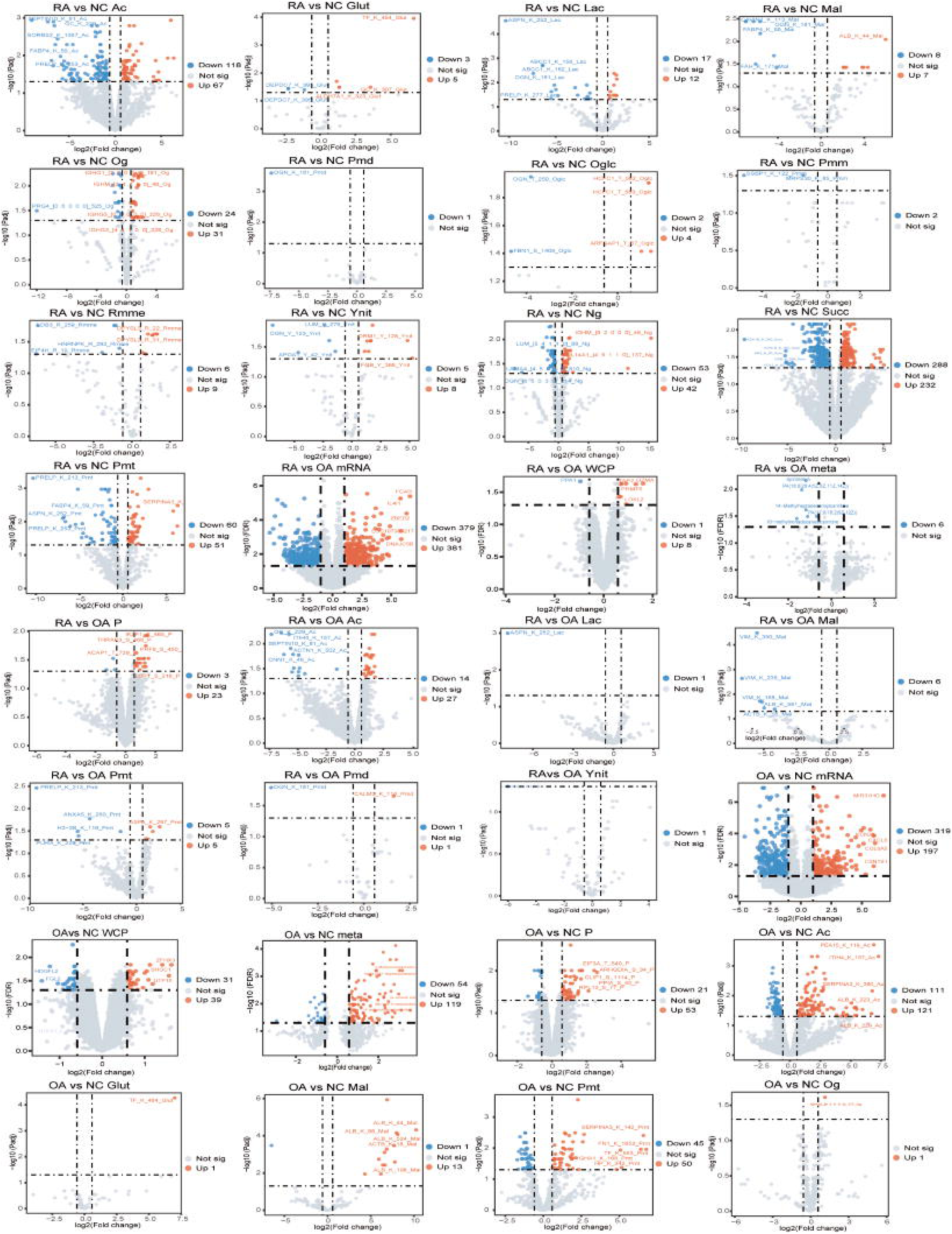
Differential analysis of quantitative results in super-omics. Volcano plots illustrate the differential analysis between each comparisons across various omics. The top five most significantly altered features are annotated.

**Figure S4.**
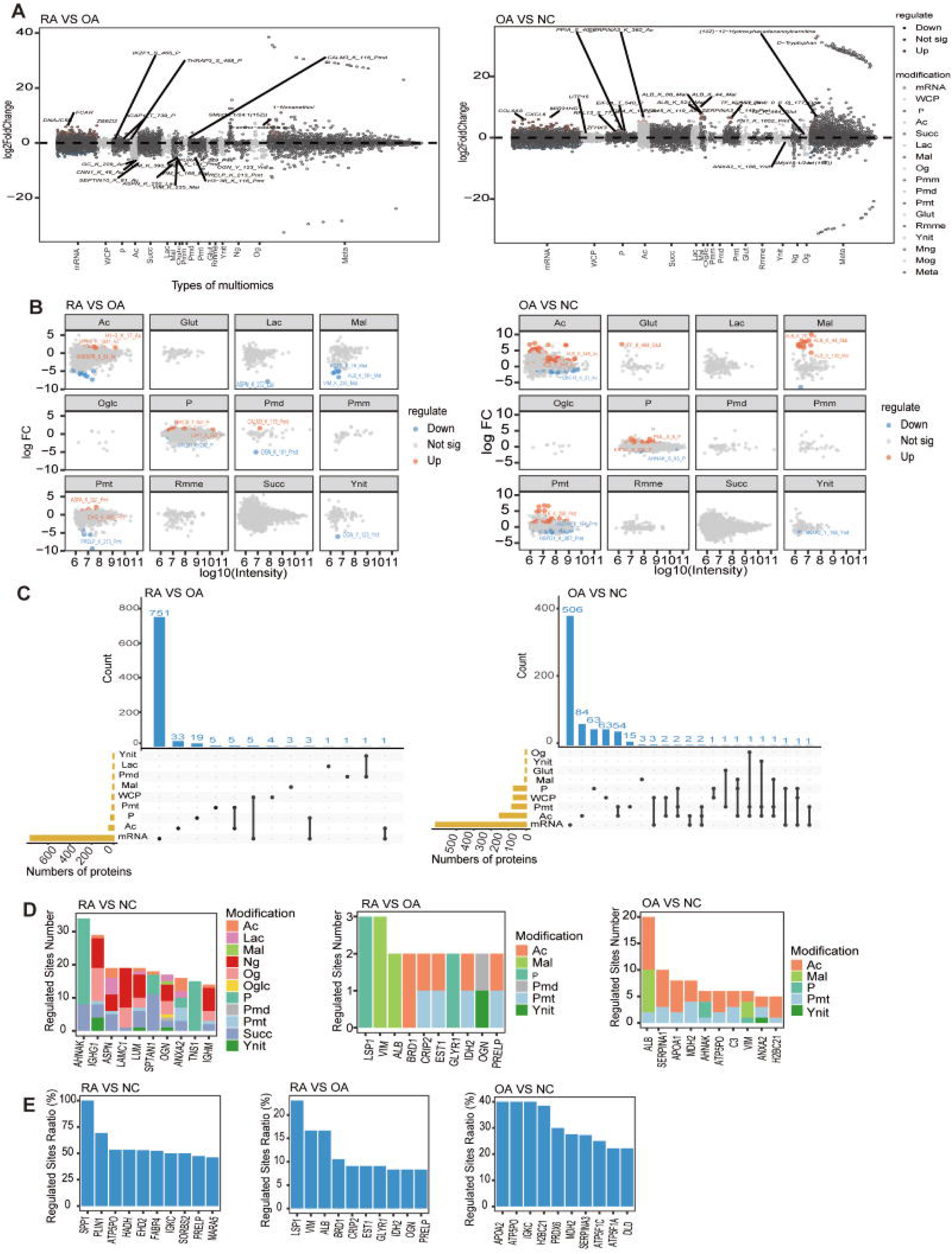
Screening of candidate clinical diagnostic biomarkers for RA. (A) Manhattan plots illustrate the overall differential analysis of transcriptomics, proteomics and 14 types of PTMs between RA and OA groups, as well as OA and NC groups. Some significantly altered entries are annotated. (B) Scatter plots illustrate the relationship between the intensity and Fold change (FC) of all detected entries across 12 types of PTMs between RA and OA groups, as well as OA and NC groups. The top three entries with the highest intensity are annotated. (C) UpSet plots illustrate the distribution of transcriptomics, proteomics, and PTMs with differential entries between RA and NC groups, as well as OA and NC groups, on the same gene. (D) Bar plots illustrate the top 10 genes with the largest number of differential PTMs sites between RA and NC, between RA and OA groups, as well as between OA and NC groups. (E) Bar plots illustrate the top genes with the largest proportion of differential PTMs sites between RA and NC, between RA and OA groups, as well as between OA and NC groups.

**Figure S5.**
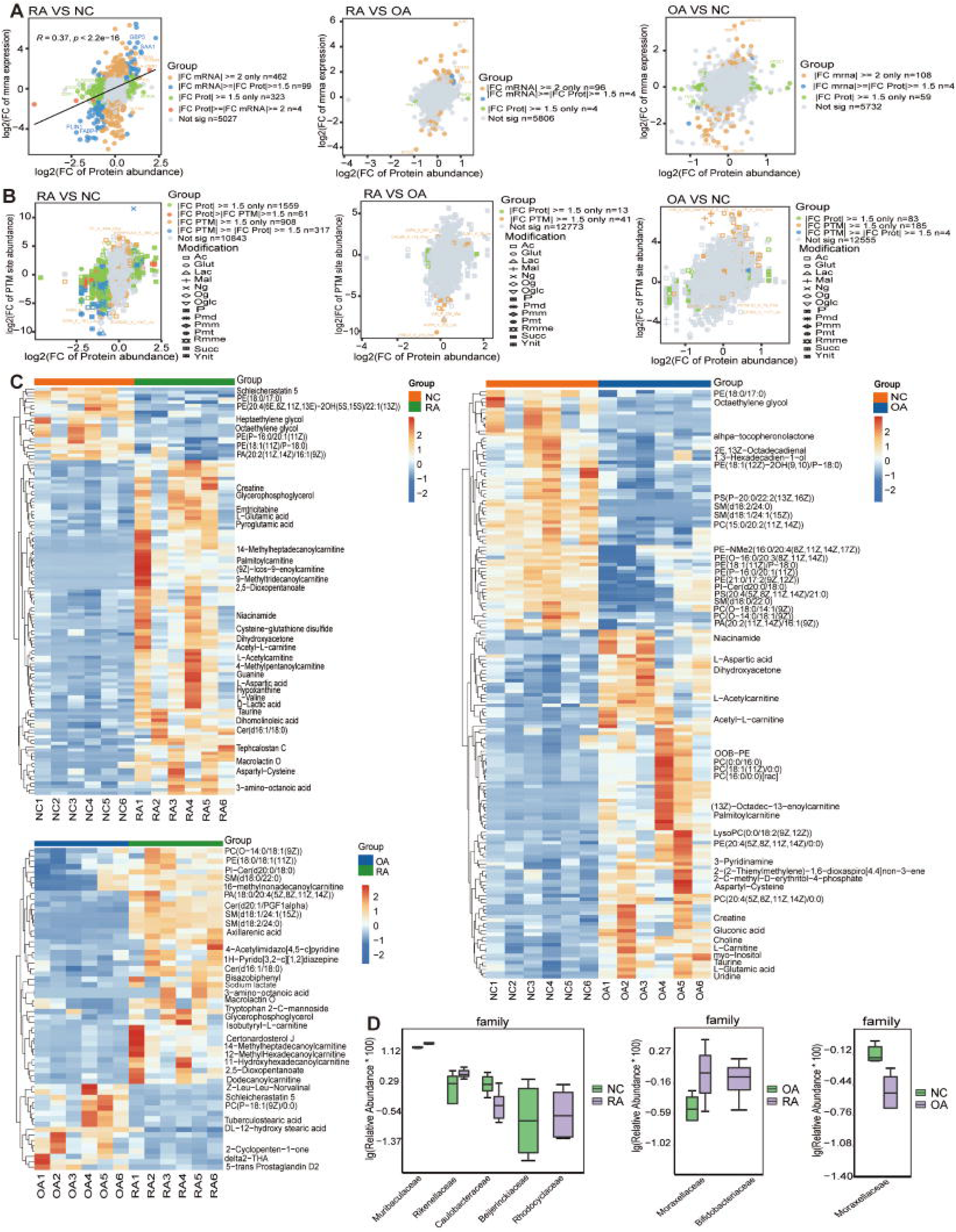
Integrated analysis between different omics. (A) Scatter plots illustrate the joint analysis of transcriptomic and proteomic data between RA and NC, between RA and OA groups, as well as between OA and NC groups. (B) Scatter plots illustrate the joint analysis of proteomic and PTMs’ data between RA and NC, between RA and OA groups, as well as between OA and NC groups. (C) Heatmap illustrate the differential metabolites between RA and NC, between RA and OA groups, as well as between OA and NC groups. (D) Bar plots illustrate the abundance of differential bacterial families between RA and NC, RA and OA groups, as well as between OA and NC groups.

**Figure S6.**
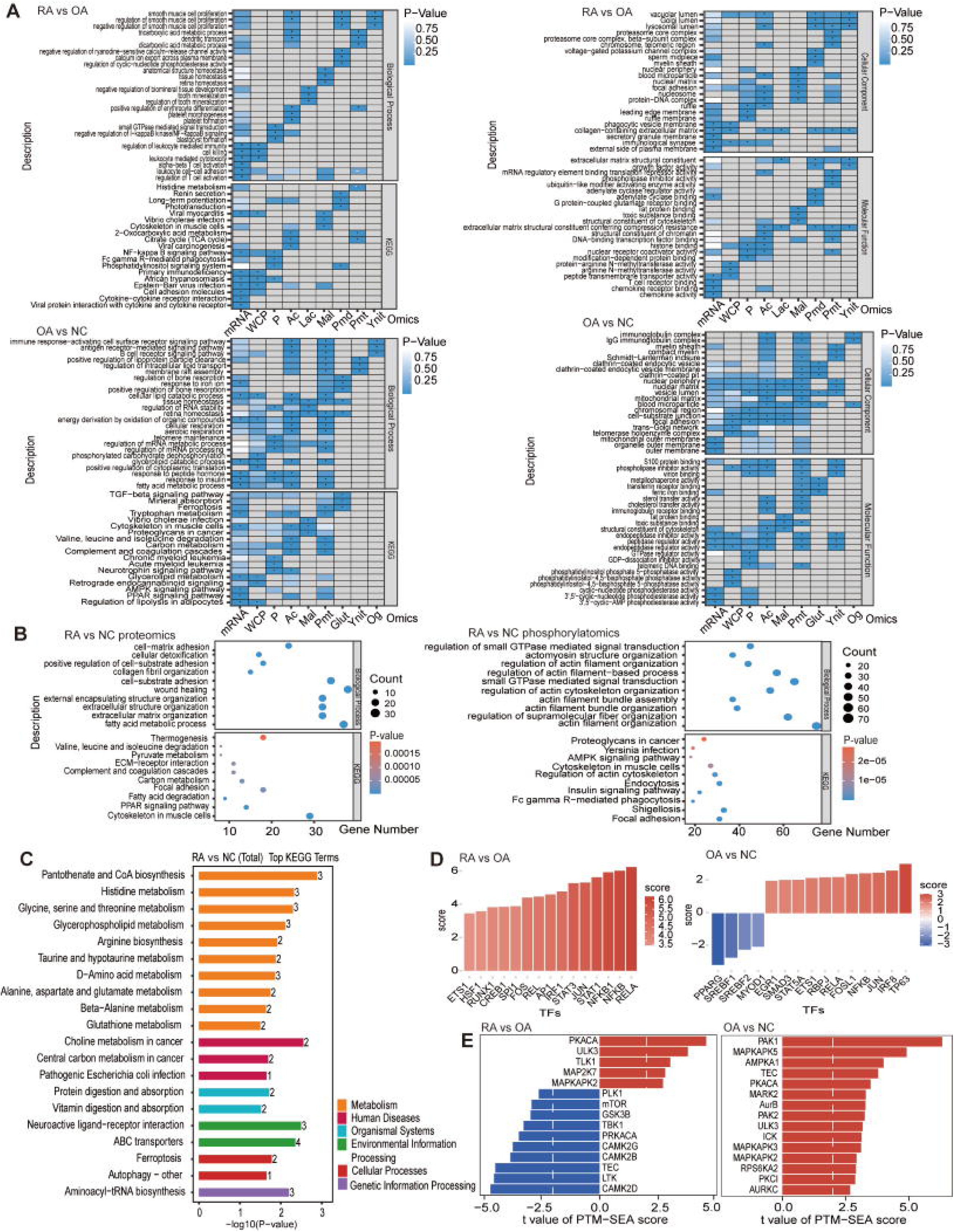
Enrichment analysis and kinase analysis of multi-omics data from different groups. (A) Heatmap illustrate Gene Ontology (GO) /Kyoto Encyclopedia of Genes and Genomes (KEGG) enrichment analysis across transcriptomics, proteomics and PTMs between RA and OA groups, as well as between OA and NC groups. *p value < 0.05, which indicates the pathway is significantly enriched. (B) Bubble plots illustrate GO/KEGG enrichment analysis for proteomics and phosphorylatomics between RA and NC groups. (C) Bar plot illustrate the enrichment analysis of differential metabolic between RA and NC groups. (D) Bar plots illustrate the predicted inflammation-related transcription factors by comparing the differential gene expression levels between RA and OA groups, as well as between OA and NC groups. (E) Bar plots illustrate the top 15 kinases identified by PTM-sea analysis between RA and OA groups, as well as between OA and NC groups.

**Figure S7.**
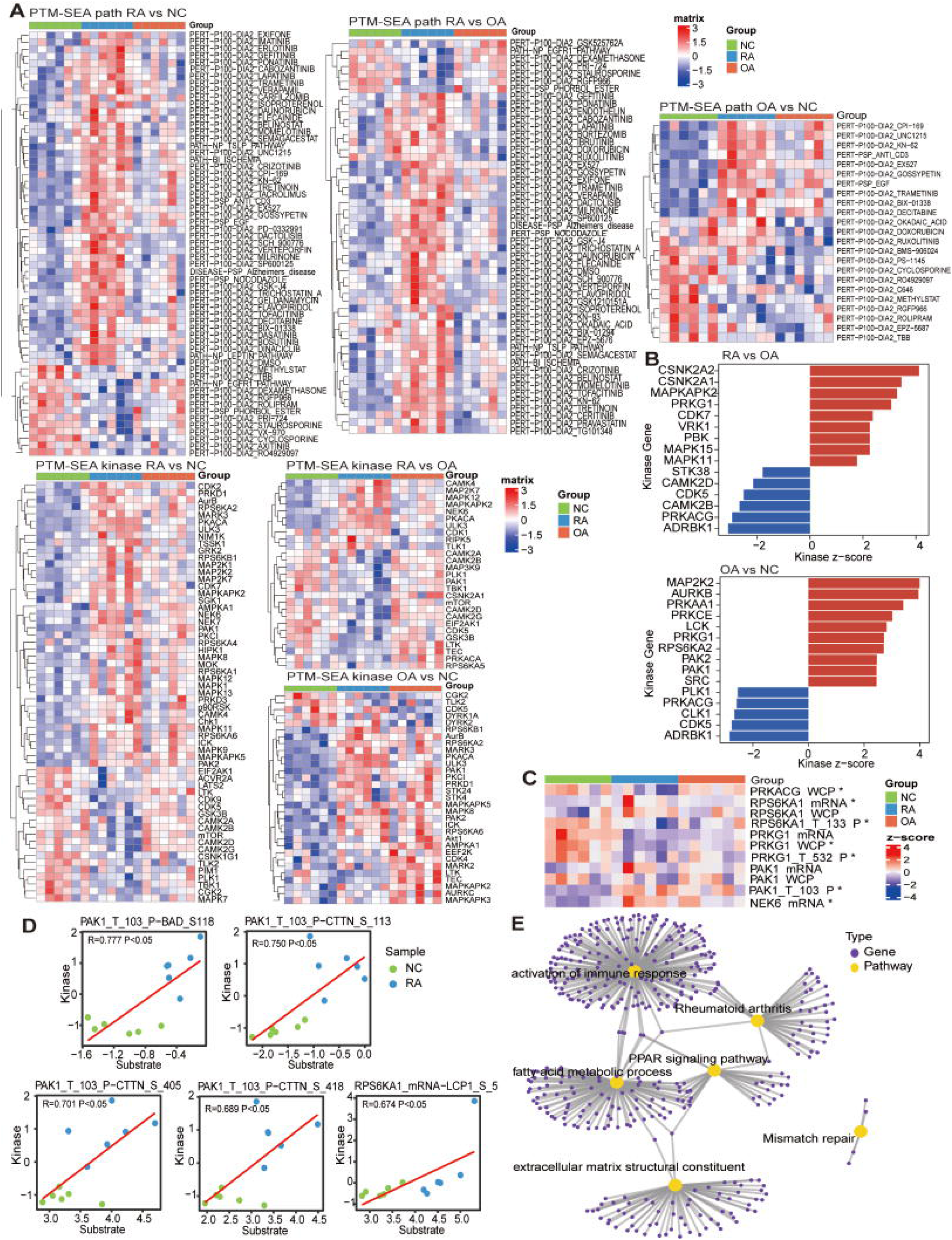
Kinase analysis of phosphorylation modifications. (A) Heatmap illustrate the PTM-SEA analysis results of pathway and kinase in RA vs NC, RA vs OA, and OA vs NC groups. (B) Bar plots illustrate the top 15 kinases identified by KSEA analysis between between RA and OA groups, as well as between OA and NC groups. (C) Heatmap illustrates the entries of transcriptomics, proteomics and differential PTMs for genes identified in Figures 4F and 4G. (D) Scatter plotS illustrate the joint analysis of kinases and their substrates between RA and NC groups. (E) The signaling pathway network diagram illustrates the relationship of differential pathways and genes with the most significant changes in RA across all omics.

**Figure S8.**
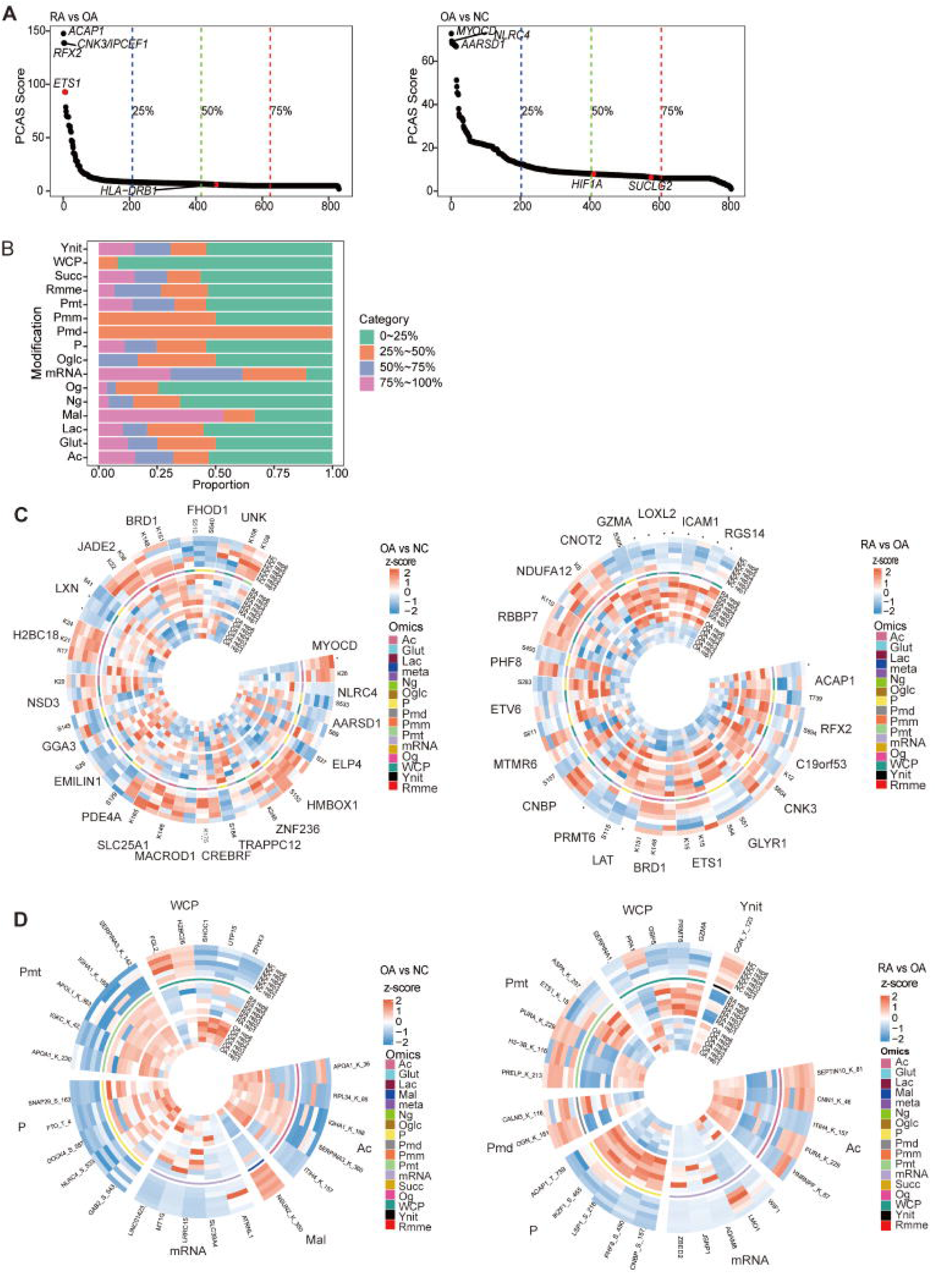
Identification of important genes in RA using PCAS analysis. (A) Scatter plots illustrate the differential genes between RA and OA groups,, as well as between OA and NC groups, ranked in descending order by PCAS score. The red dots indicate RA-related genes that have been previously reported. (B) Stacked bar plot illustrates the proportion of differential entries associated with genes based on quartile PCAS score between RA and NC groups across each omics relative to the total number of differential entries for all genes based on PCAS score within the same omics dataset. (C) Circos plots illustrate z-score intensity of top 20 genes based on PCAS score between RA and OA groups, as well as between OA and NC groups, across transcriptomics, proteomics and differential PTMs. *p value < 0.05, which indicates differential transcriptomics or proteomics entries. (D) Circos plots illustrate the z-score intensity of 5 genes with the highest absolute values of FC in each omics from top 25% differential genes based on PCAS score between RA and OA groups, as well as between OA and NC groups.

**Figure S9.** Upstream and downstream regulatory genes of high PCAS score genes and their changes. The diagrams illustrate upstream and downstream relationships of PLIN1, LDB3, ITIH3, RGCC, and ADH1C, PLIN1, LDB3, ITIH3, RGCC, and ADH1C. The diagrams show the alterations in PLIN1, LDB3, ITIH3, RGCC, and ADH1C, and corresponding changes in their upstream and downstream genes.

**Figure S10.** Network analysis and enrichment analysis of PCAS results. (A) Network diagrams for Modules 1, 2, 3, 5, 6, and 7. The top 25% genes based on PCAS score between RA and NC groups were grouped into seven modules. The Network diagram showing three interactions between genes such as INTACT, PTM and TF, as well as the value of the gene’s PCAS score. (B) Heatmap illustrate the GO/KEGG pathway analysis for quartile genes based on PCAS score between RA and OA groups, as well as between OA and NC groups.

**Figure S11.** The network and pathway joint analysis of the PCAS results. (A) Scatter plots illustrate the genes related with “Cell-Substrate Junction,” “Integrin-Mediated Signaling Pathway”, “Fatty Acid Metabolism”, “Pyruvate Metabolism”, and “MYC Targets v1” pathways among quartile genes based on PCAS score between RA and NC groups. The red dots indicates the top 3 genes with the highest PCAS scores in the pathway. (B) Network diagrams for Module 5, 6 and important pathways. The Network diagram illustrates three interactions between genes such as INTACT, PTM, and TF. Meanwhile, diagrams also illustrate the value of the gene PCAS score, and the signaling pathway. (C) Heatmap illustrates the correlation between the most functionally important genes in Figure 6F and clinical indicators. CRP, C-reactive protein; ESR, erythrocyte sedimentation rate; RF, rheumatoid factor; DAS28, Disease Activity Score-28; CCP, Anti-cyclic citrullinated peptide. “SEX” is coded as 1 for male and 0 for female. (D) The FLSs transfected with small interfering RNA (siRNA) targeting STK17B for 24 hours, then stimulated with TNF-α for 24 hours. ***p<0.001. (E) The FLSs stimulated with TNFα (10ng/ml), along with DRAK2-IN-1 (1μM) and Quercetin (80μM) for 24h, respectively. *p<0.05, **p<0.01,***p<0.001.

## Notes

### Competing Interest Statement

The authors have declared no competing interest.

### Summary of Updates

Corrections have been made to some errors in the figure, results, and discussion.

